# A benchmark study of protein folding algorithms on nanobodies

**DOI:** 10.1101/2022.08.07.503071

**Authors:** Shibo Liang, Ziquan Liang, Zecheng Wu, Feijuan Huang, Xu Wang, Zihao Tan, Rui He, Zeyi Lu, Yuanzhe Cai, Bingding Huang, Xin Wang

## Abstract

Nanobodies, also known as single domain or VHH antibodies, are the artificial recombinant variable domains of heavy-chain-only antibodies. Nanobodies have many unique properties, including small size, good solubility, superior stability, rapid clearance from blood, and deep tissue penetration. Therefore, nanobodies have emerged as promising tools for diagnosing and treating diseases. In recent years, many deep-learning-based protein structure prediction methods have emerged that require only protein sequences as input to obtain highly-credible 3D protein structures. Among them, AlphaFold2, RoseTTAFold, DeepAb, NanoNet, and tFold performed excellently in protein prediction or antibody/nanobody prediction. In this study, we selected 60 nanobody samples with known experimental 3D structures in the Protein Data Bank (PDB). Next, we predicted their 3D structures using these five prediction algorithms from only their 2D amino acid sequences. Then, we individually compared the predicted and experimental structures. Finally, the results are analyzed and discussed.

## Introduction

Proteins comprise polypeptide chains of dehydrated and condensed amino acids, which are then coiled and folded to form large, spatially structured biomolecules. The 3D structure of proteins is essential for understanding their biological functions. Understanding how amino acid sequences determine the 3D protein structure is a major challenge in the field, also known as the “protein folding problem”.^1^

Antibodies (Abs) are an extensively studied class of proteins due to their important functions. Abs, also known as immunoglobulins (Ig), are large Y-shaped proteins secreted mainly by plasma cells. Abs comprises four polypeptide chains linked by interchain disulfide bonds in most mammals, including humans. The two chains with larger molecular weights are called heavy (H) chains, and the two chains with smaller molecular weights are called light (L) chains. The H and L chains consist of a relatively stable constant I region and a more variable (V) region. Abs can also be divided into two antigen-binding fragments (Fabs), containing one VL, VH, CL, and CH domain, and the crystallizable fragment (Fc).^2^

The V regions of the H and L chains are called VH and VL, respectively. Each has three highly variable amino acid regions called complementarity determining regions (CDRs), CDR1, CDR2, and CDR3, of which CDR3 is the most variable. The three CDRs of VH and VL together form the Ig antigen-binding site, which determines the AB’s specificity and is the site where it recognizes and binds to the antigen.^3,4^ Outside of the three CDR regions, the composition and order of amino acids in the V region are relatively conserved, comprising four framework regions (FRs), FR1, FR2, FR3, and FR4, for each VH or VL.

Belgian scientists discovered a new Ab in the serum of camelids in 1993, which were later also identified in dromedaries and alpacas.^5^ The new Ab’s structure was missing the L chain C region, H chain C region (CH1), and two L chain V regions. It was only composed of two H chain V regions and H chain C regions (CH2 and CH3). The structure of this Ab differs from those of conventional Abs, leading it to be called a heavy-chain Ab.^5,6^

The V domain of H chain Abs (VHH)^7^ has a molecular mass based on in vitro recombinant protein expression of ~15 kDa, about one-tenth that of conventional Abs. The VHH retains the AB’s total antigen-binding capacity and is known as a nanobody (Nb).^8^

Nbs have several advantages over conventional Ab Fab and single chain Ab fragments, including weaker immunogenicity, lower production cost, greater water solubility, and superior tissue permeability and stability.^8,9^ These properties enable Nbs to access sites inaccessible to conventional Abs, such as inside tumor cells^10^ and the blood-brain barrier,^11^ enabling their use in more applications.

Currently, protein structures are determined experimentally by expensive and time-consuming techniques such as X-ray crystallography,^12^ cryo-electron microscopy (cryo-EM),^13^ and nuclear magnetic resonance.^14^ Over the last 60 years, these labor-intensive efforts have determined the structures of ~170,000 of the 200 million known proteins in all organisms.^15^ The development of a method that can accurately deduce protein structure from amino acid sequences alone would greatly benefit many biomedical research fields, such as Ab drug discovery.

Deep-learning-based protein structure prediction algorithms have emerged in recent years that can predict 3D protein structures from their 2D protein sequences. Among them, AlphaFold2 (AF2) was the first to reach relatively high accuracy.^16^ RoseTTAFold,^17^ which is open-source and was released almost simultaneously with AF2, also has a high prediction ability. In addition, the DeepAb, NanoNet, and tFold platforms are also reported to show good performance in protein and Ab/Nb prediction.

AF2^16^ is a deep-learning-based program developed by the Google DeepMind Team that won the protein structures competition at Critical Assessment of Structure Prediction (CASP) 14 in November 2020. Differing from experimentally obtained structures by only one atom’s width on average, AF2 reached a comparable level of protein structure prediction to humans using sophisticated instruments such as cryo-EM. Compared to AlphaFold1 (AF1), developed in 2018, AF2 overcame AF1’s limitation that it could only predict protein structures in the CASP 13 dataset.^18^ Importantly, protein structures predicted by AF2 were also much more accurate than those by AF1.^19(p13)^

AF2’s algorithm can be divided into feature extraction, encoding, and decoding. Once provided with a protein’s amino acid sequence, AF2 searches the database for homologous sequences to obtain a feature representation of the sequence and amino acid pairs. Next, the encoder constructs a multiple sequence alignment (MSA)^20^ and pair matrices, and the information in the two matrices is updated. Finally, the encoder encodes the protein structure’s 3D coordinates using relative positions.

AF2 has four main features. First, the database used for neural network training is very large, including a 1.7 TB big fantastic database (BFD) built by AF2’s authors and the Uniref90, Mgnify, and Uniclust30 databases,^21–24^ greatly improving its prediction model’s accuracy. Second, the whole Evoformer model and structure module both use recycling, where the output is rejoined with the input for information refinement. Third, AF2 extensively uses the transformer structure,^25^ and its MSA and amino acid pair information are updated using the attention mechanism. Finally, a bidirectional encoder representation from transformers-like masking operation^26^ adds noise to the various input information to ensure stable output, improving robustness and generalization.^27^

Huge databases support AF2. AF2 uses multiple sequence databases, of which BFD is the biggest. BFD is one of the largest publicly available collections of protein families (https://bfd.mmseqs.com/), comprising 2,204,359,010 protein sequences from 65,983,866 families sourced from reference databases, metagenomes, and metatranscriptomes and represented as MSAs and hidden Markov models (HMMs).

Different steps in the AF2 algorithm use distinct databases. When searching for template structures for prediction, AF2 uses the PDB^28^ and PDB70^21^ clustering databases. PDB70 contains HMM profiles on a representative set of PDB sequences, filtered with a maximum pairwise sequence identity of 70%. It currently contains 35,000 HMM profiles with an average length of 234 matched states.

AF2’s MSA lookup during training and prediction uses the Uniref90^22^ (v.2020_01), BFD, Uniclust30^23^ (v.2018_08), and Mgnify^24^ (v.2018_12) databases. The Uniref database (UniProt reference clusters) provides a set of sequence clusters from the UniProt knowledge base and selected UniParc records to completely cover the sequence space at multiple resolutions (100%, 90%, and 50% identity) while hiding redundant sequences. Uniref90 is created by clustering Uniref100 sequences at the 90% sequence identity level. The Uniclust30 database provides functionally homogeneous sequence clusters at 30% sequence identity clustering depths, representative sequence sets, cluster MSAs, and sequence annotations using Pfam, Structural Classification of Proteins, and PDB matches.

Mgnify is a free-to-use platform for assembling, analyzing, and archiving microbiome data obtained by sequencing microbial populations in particular environments. AF2 uses Uniclust30 (v.2018_08) to construct a distillation structure dataset.

RoseTTAFold,^17^ first published online in July 2021, was developed by David Baker’s team at the University of Washington. Althoguh RoseTTAFold’s network architecture is generally similar to AF2, it claims a comparable performance through a different three-track network in which information at the 1D sequence level, 2D distance map level, and 3D coordinate level are sequentially transformed and integrated. The databases used by RoseTTAFold also differ from AF2. While AF2 uses the BFD (1.7 TB), PDB70 (56 GB), Uniref90 (58 GB), and MGnify (64 GB), params (3.5 GB), small_BFD (17 GB), PDB_mmcif (206 GB) and Uniclust30 (24.9 GB) databases (total size of 2.2TB), RoseTTAFold uses BFD (1.7 TB), PDB100 (666 GB), and Uniref30 (194 GB) databases (total size of 2.6TB).^16,17^ Nevertheless, the size of databases used by both methods is also comparable.^29^

RoseTTAFold extends the AF2 two-track architecture to a third track, operating in three-dimensional coordinates to provide a tighter association between sequence, residue-residue distances/orientations, and atomic coordinates. In this architecture, information flows back and forth between 1D amino acid sequence information, 2D distance metrics, and 3D coordinates, enabling the network to jointly determine relationships, distances, and coordinates within and between sequences. In contrast, the AF2 architecture only infers 3D structural coordinates after the 1D and 2D information is processed by the neuron network. This network also enables the rapid generation of accurate protein-protein complex models from sequence information alone, surpassing traditional approaches that model individual subunits before docking.

While RoseTTAFold’s innovative three-track model does not perform as well as AF2 on the CASP 14 data due to training database and hardware issues, it still shows relatively high predictive accuracy. In addition, RoseTTAFold takes less time and computer resources to produce predictions than AF2, which is a significant benefit to RoseTTAFold beside it fully open-source characteristics.

DeepAb, first published online in June 2021, was developed by Jeffrey A. Ruffolo’s team at Johns Hopkins University. Its algorithm has two phases. The first phase is a deep residual convolutional network (ResNet)^30^ and a long and short-term memory network (LSTM)^31^ that specifically predicts the relative distance and direction (2D structure) between residual pairs in the V region (Fv). The second phase is a distance and angle-constrained approach based on the Rosetta minimization protocol for predicting 3D structures. The network requires only H and L chain sequences as input and is designed with interpretable attention components to provide insight into the model’s prediction regions.^32^

DeepAb was found to perform better than its previous iteration, DeepH3,^33^ on the testing set of therapeutically relevant Abs and the RosettaAb benchmark set.^34^ The DeepAb framework is based on the CDR H3 ring structure prediction method. However, it enhances it with two innovations. First, DeepAb enriches Ig sequences using supervised learning in the Fv region with a recurrent neural network (RNN) model. The RNN embeds representations of general feature patterns in the Ig sequences, with a fixed implicit layer size of 64, allowing training on features that are not evident in a small subset of known structures. The encoder in the RNN model uses a bidirectional LSTM network (Bi-LSTM), and the decoder uses an LSTM.^31^ Second, DeepAb represents the Fv structure as a set of inter-residue distances and orientations, predicting inter-residue distances between three atom pairs and the set of inter-residue dihedrals and planar angles.^32^

NanoNet was developed by Tomer Cohen’s team at the Hebrew University of Jerusalem and is the first modeling approach optimized for VHH. The model is constructed using a large amount of data to train the neural network using the Ab’s VH domain, the Vβ domain of the T cell receptor (TCR), and VHH ^35^.

While its primary goal is accurate Nb modeling, NanoNet can also accurately model the VH domains of Abs and the Vβ domains of TCRs since they are included in the training dataset. NanoNet’s training dataset uses abYbank/AbDb^36^ and SabDab^37^, comprising 2,085 total non-redundant structures of Nbs (319) and monoclonal Ab (mAb) H chains (1,766) were used. Its deep learning model accepts amino acid sequences (Nb, mAb VH domains, or TCR Vβ domains) as input and generates alpha carbon (Cα) atom coordinates. Like AF2 and RoseTTAFold, NanoNet also used direct end-to-end learning, allowing the network to learn complete 3D structures without splitting the modeling problem into frame/CDR modeling.^35^ This unique model that focuses only on Cα atomic coordinates allows it to predict Ab results with much better accuracy and significantly reduced computation time.

Tencent artificial intelligence (AI) Lab’s iDrug is an AI-driven open platform for pre-clinical drug discovery and development based on their self-developed deep learning algorithms with database and cloud computing support. The primary iDrug component focusing on protein folding is tFold. Tencent AI Lab reported using tFold to help resolve the crystal structure of steroid 5 alpha-reductase 2 on November 17, 2020. It effectively improved the accuracy of protein structure prediction through their tFold algorithm.^38,39^

The tFold algorithm uses three innovative techniques to improve modeling accuracy. First, they developed a multi-source fusion technology to mine the co-evolutionary MSA information. Then, with the help of a deep cross-attention residual network, tFold can greatly improve the prediction of important 2D structural information, such as residue-residue distances and orientation matrices. Finally, a novel template-based free modeling method is used to integrate the structural information in the 3D models generated by free modeling and template-based modeling, greatly improving the accuracy of the final 3D model.^39^

To accurately and visually benchmark these five protein prediction algorithms for Nb 3D structures, we selected two popular methods for calculating and analyzing their strengths and weaknesses: root mean squared differences (RMSDs; Fig. 1A) and template modeling I scores between the experimental (gold standard) and prediction results. We also generated Rasch plots for each sample using the five algorithms in the supplemental files.

**FIGURE 1.**
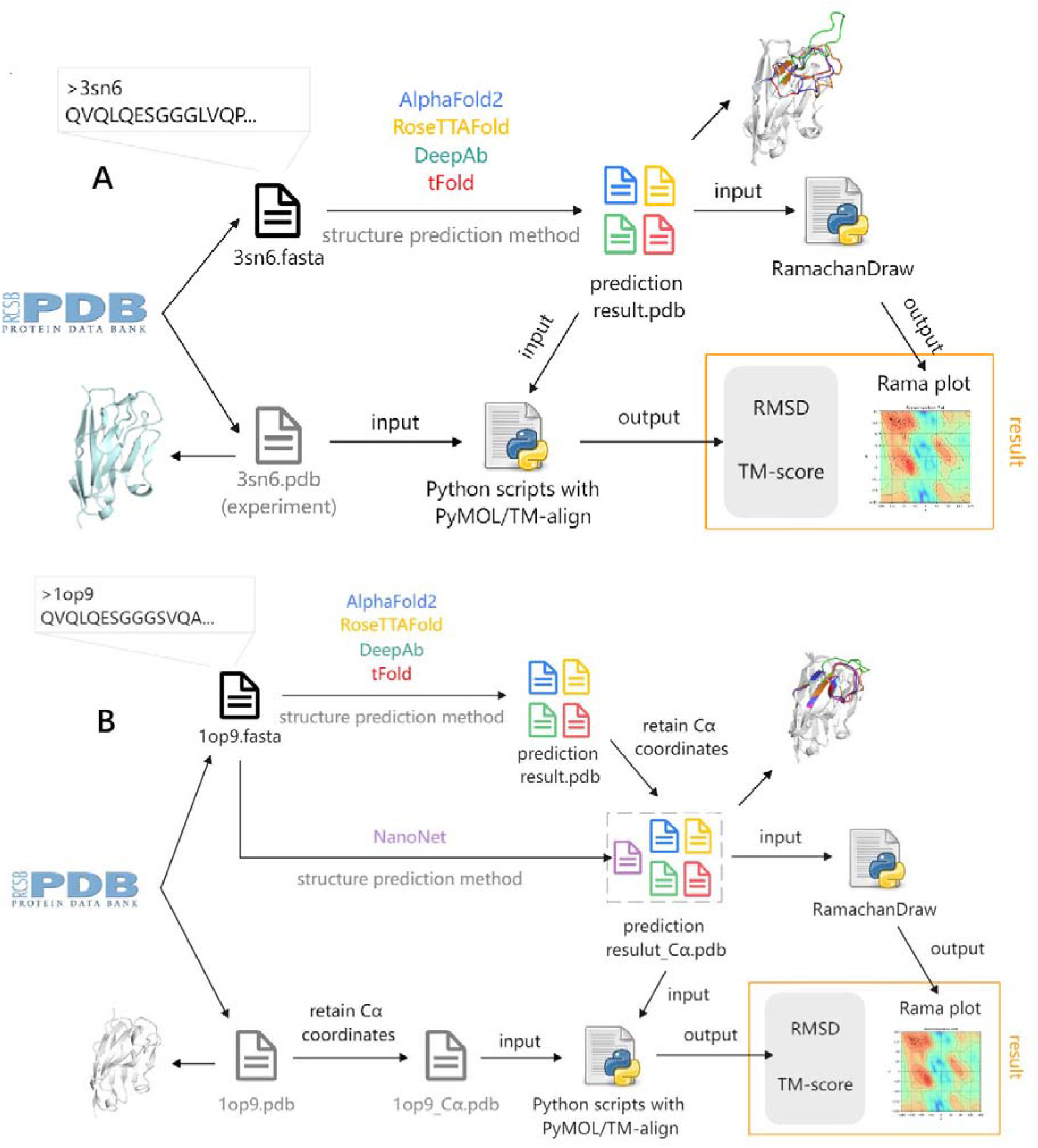
A schematic diagram of the two comparisons. (A) Schematic of how scores were calculated for all-atom protein models. (B) Schematic of how scores were calculated for Cα-only protein models.

RMSD is a parameter used to represent the difference in atomic positions between protein structures.^40^ It should be noted that the RMSD calculation requires identical atomic order and number in the two protein structure files. However, NanoNet’s output is a protein structure file of only Cα atomic coordinates. This difference creates the problem of unequal atom numbers when comparing it to the experimental file of all atomic coordinates. Therefore, we deleted the non-Cα atomic coordinates from the experimental file when calculating each NanoNet model’s RMSD.

To include NanoNet in the comparison of RMSD differences among the five methods and the experimental file, we created copies of the protein structure files containing only Cα atomic coordinates from the full atomic protein structure prediction files of the other four methods. We then calculated their Cα-only RMSDs using these Cα-only files (Fig. 1B).

The TM score is a measure of similarity between two protein structures.^41^ The TM score was designed to be a more accurate measure of overall similarity to a full-length protein structure than the commonly used RMSD measure. The Rasch plot is a method for visualizing the dihedral angles ψ and φ of amino acids in the main chain of a protein structure. The Rasch plot enables an assessment of whether the protein’s conformation is plausible or not.^42^

To compare the performance of the five algorithms on Nb protein structure prediction, we selected 60 samples with experimentally determined 3D structures in the PDB database. The five prediction methods described above were then run on their input 2D protein sequences. Based on this large set of 3D atomic coordinate prediction data, we compare the prediction results of the five algorithms with the experimental data to obtain RMSDs and TM scores. We then analyze and discuss our results.

## Results

A scheme of how we compared the all-atom protein models with four of the five protein folding prediction methods is shown in Figure 1A. NanoNet was excluded since it only predicts Cα coordinate models. First, we searched for suitable Nbs in the PDB database and downloaded their protein sequence (FASTA format) and structure (PDB format) files. Second, we imported the protein sequence files into AF2, RoseTTAFold, DeepAb, and tFold to obtain protein structure prediction results as PDB format files. Third, these protein structure prediction results were imported into the RamachanDraw script to obtain a Ramachandran plot. Finally, the experimental and predicted protein structure files were imported into the PyMOL script to calculate the RMSDs and TM scores for these four methods.

The worst RMSDs were obtained with DeepAb (Fig. 2A; Supplementary Table 1) with a mean VHH RMSD of 4.19±1.39 Å (Table 1). The mean RMSDs of the four methods did not differ significantly in the FR. The FR spatial structure can be predicted more accurately by all four methods compared to CDRs due to its relative stability (Table 1).

**FIGURE 2:**
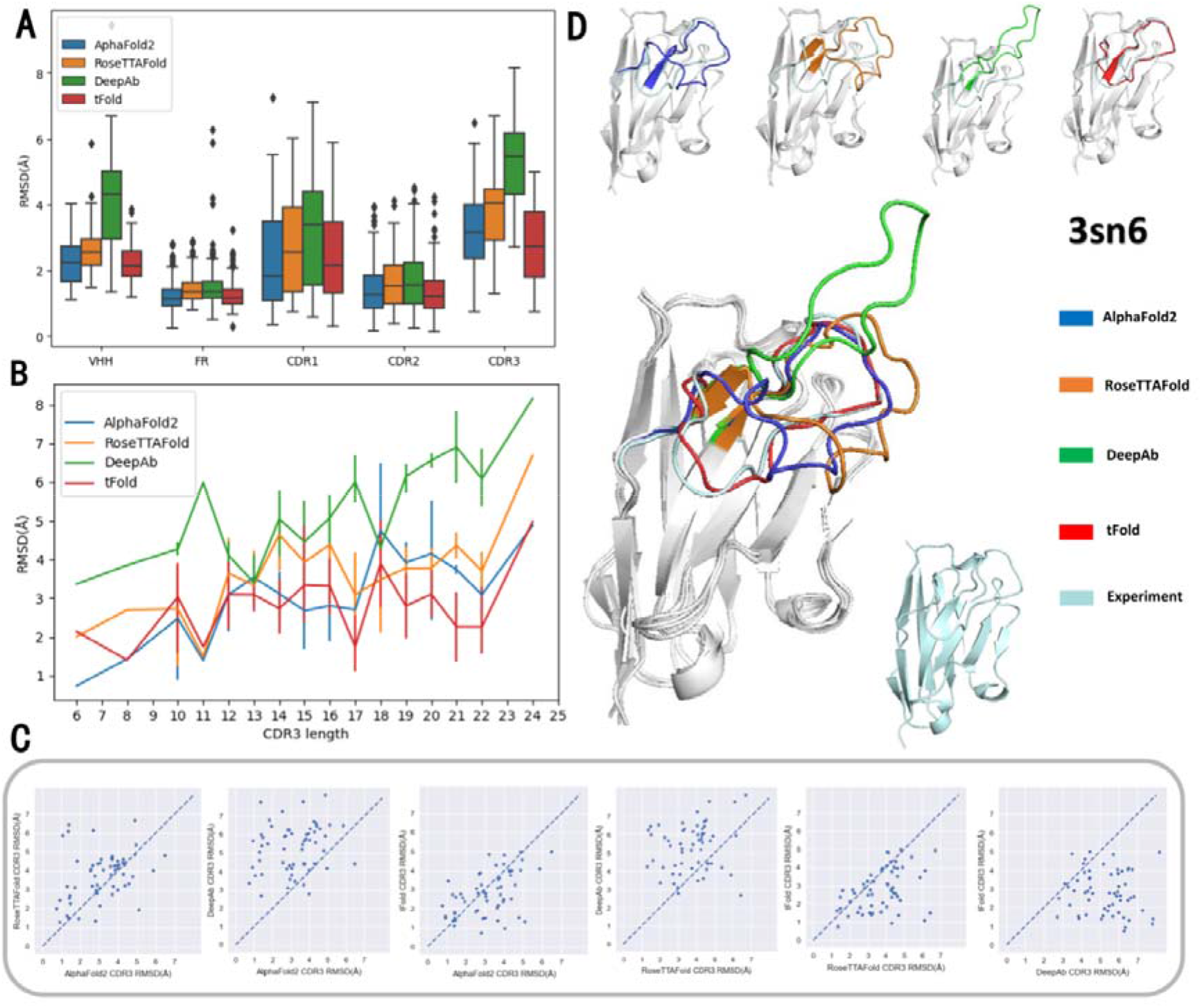
RMSDs and structure comparison of the four all-atom prediction methods with the experimental structure. (A) Boxplot of RMSDs for the full VHH structure, FR, CDR1, CDR2, and CDR3. (B) Length comparison of CDR3 loop RMSDs in the line graph. (C) Scatterplots of pairwise RMSD comparisons for CDR3 loops among the four different methods. (D) Visual comparison of the predicted 3D Nb structures in PyMOL.

**TABLE 1:** Mean RMSDs between experimental and all-atom predicted whole VHH and FR, CDR1, CDR2, and CDR3 structures for each method.

| Method | VHH | FR | CDR1 | CDR2 | CDR3 |
| --- | --- | --- | --- | --- | --- |
| AlphaFold2 | 2.25($\pm 0.69$) | 1.19($\pm 0.42$) | 2.35($\pm 1.54$) | 1.52($\pm 0.99$) | 3.17($\pm 1.32$) |
| RoseTTAFold | 2.59($\pm 0.77$) | 1.42($\pm 0.40$) | 2.80($\pm 1.52$) | 1.69($\pm 0.95$) | 3.74($\pm 1.27$) |
| DeepAb | 4.19($\pm 1.39$) | 1.48($\pm 0.64$) | 3.14($\pm 1.60$) | 1.78($\pm 1.05$) | 5.28($\pm 1.28$) |
| tFold | 2.20($\pm 0.59$) | 1.25($\pm 0.39$) | 2.44($\pm 1.35$) | 1.45($\pm 0.96$) | 2.79($\pm 1.14$) |

AF2 showed the best results in the CDR1 loop region with a mean RMSD of 2.35±1.54 Å, followed by tFold with a mean RMSD of 2.44±1.35 Å (Table 1). However, a relatively high outlier (3K3Q, with a CDR1 loop length of 19) of 7.25 Å was seen with AF2 (Supplementary Figure 1). The worst performing method was DeepAb with a mean RMSD of 3.14±1.60 Å, followed by RoseTTAFold with a mean RMSD of 2.80±1.52 Å (Table 1).

All methods had a relatively low RMSD value in the CDR2, which did not differ significantly among them (Fig. 2A and Table 1). In addition, average CDR1, CDR2, and CDR3 lengths in our test dataset were the smallest for CDR2 (Supplementary Figure 2 and Supplementary Table 4).

Large RMSD differences begin to appear between the four methods in the CDR3, which is more challenging to predict than CDR1 and CDR2 due to its highly variable nature and the longer residue lengths. The best prediction was by tFold with a mean RMSD of 2.79±1.14 Å, followed by AF2 was a mean RMSD of 3.17±1.32 Å. The worst performing method was DeepAb with a mean RMSD of 5.28±1.28 Å, followed by RoseTTAFold with a mean RMSD of 3.74±1.27 Å (Table 1).

We further studied the relationship between CDR length and RMSD by comparing CDR3 loop length with the resulting RMSDs (Fig. B). There was a positive correlation between CDR3 length and RMSD, where RMSDs increased with CDR3 loop length and showed a sawtooth-like increase with the best performing tFold method (smallest RMSDs) and the worst performing DeepAb method (largest RMSDs).

We also performed one-to-one comparisons of RMSDs for CDR3 loops among the four methods (Fig. 2C). The comparison of AF2 with RoseTTAFold showed that points were mainly clustered around the diagonal. However, AF2 had slightly better results than RoseTTAFold overall. The comparison of AF2 with DeepAb showed that AF2 performed significantly better than DeepAb. The comparison of AF2 with tFold showed that tFold performed slightly better than AF2. The comparison of RoseTTAFold with DeepAb showed that RoseTTAFold performed significantly better than DeepAb. The comparison of RoseTTAFold with tFold showed that tFold performed much better than RoseTTAFold. The comparison of DeepAb with tFold showed that tFold performed significantly better than DeepAb.

We also visualized the structure of the prediction results of each method using PyMOL. The CDR3 loop region for which DeepAb’s prediction differed greatly from the experimental structure is highlighted in Figure 2D. In contrast, the tFold predicted structure for the CDR3 loop region was broadly consistent with the experimental structure.

A scheme of how we performed the Cα-only protein model comparison with the five protein folding prediction methods is shown in Figure 1B. First, we searched for suitable Nbs in the PDB database and downloaded their protein sequence (FASTA format) and structure (PDB format) files. Second, the protein sequence files were imported into AF2, RoseTTAFold, DeepAb, and tFold to obtain protein structure prediction results. Third, these all-atom prediction files were filtered to retain only Cα’s coordinates, enabling their comparison with NanoNet. Fourth, the protein sequence files were imported into NanoNet to obtain Cα-only protein structure prediction results. Fifth, the protein structure prediction results were imported into the RamachanDraw script to obtain a Ramachandran plot. Finally, the experimental and predicted protein structure files were imported into the PyMOL script to calculate RMSDs and TM scores for the five methods.

Comparing boxplots of Cα-only RMSDs for all methods (Figure 3A and Supplementary Table 2) showed that DeepAb performed the worst, with AF2, tFold, and NanoNet all showing superior results. NanoNet performed best on the full VHH structure but also had more outliers. The five methods performed approximately the same in the FR, but there were also outliers, especially for the DeepAb and NanoNet methods.

**FIGURE 3:**
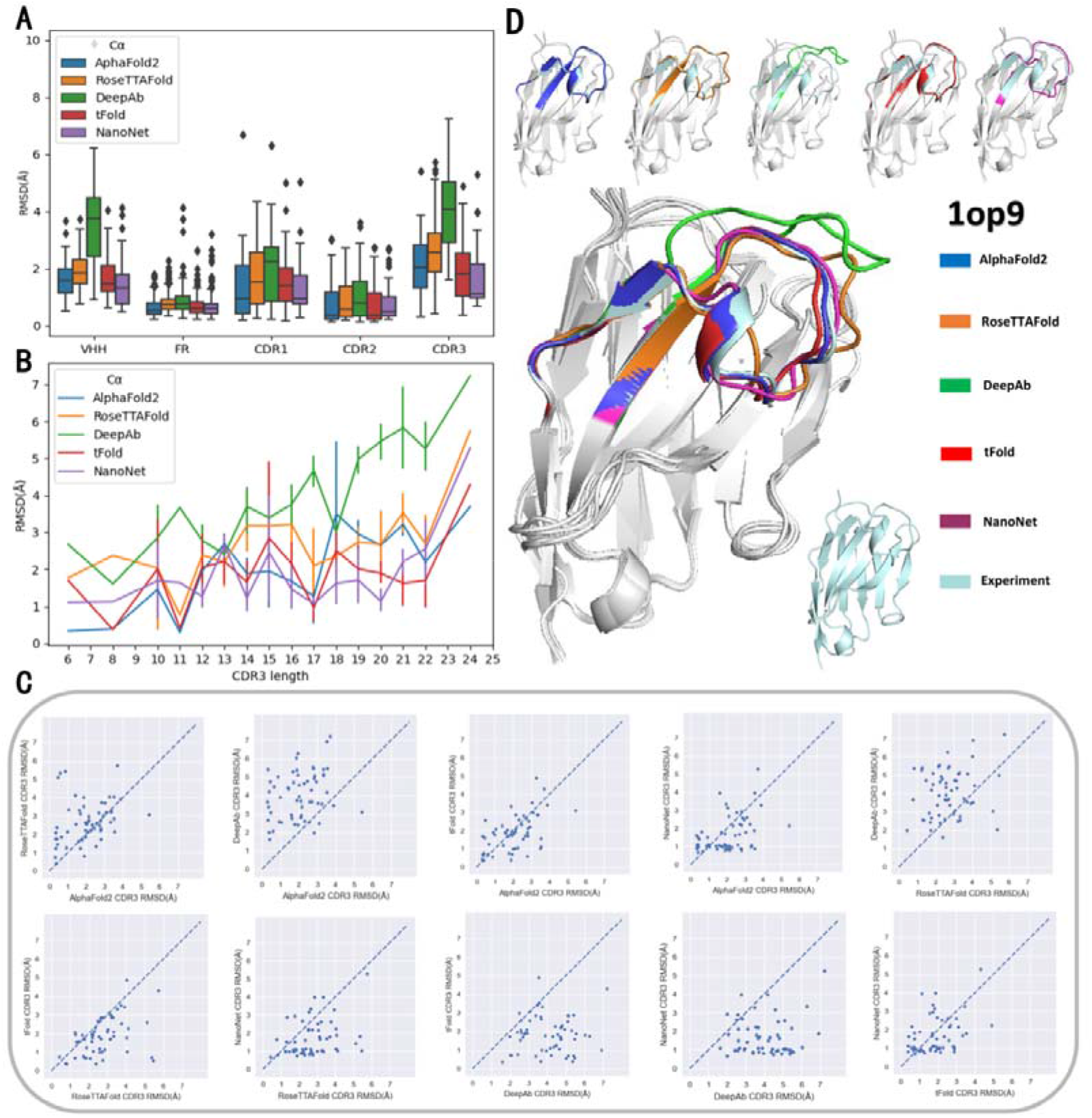
RMSD and Cα-only structure comparison of the five methods with the experimental structure. (A) Boxplot of RMSDs for the full VHH structure and FR, CDR1, CDR2, and CDR3. (B) Length comparison of CDR3 loops RMSDs in the line graph. (C) Scatterplots of pairwise RMSD comparisons for CDR3 loops among the five different methods. (D) Visual comparison of the predicted Nb 3D structures in PyMOL.

NanoNet performed best on the CDR1 ring region (Fig 3A) with a mean RMSD of Å, followed by AF2 with a mean RMSD of 1.37±1.16 Å (Table 2). The worst performing method was still DeepAb with a mean RMSD of 1.96±1.16 Å, followed by RoseTTAFold with a mean RMSD of 1.72±1.03 Å. The middle-performing method was tFold with a mean RMSD of 1.58±0.97 Å.

**TABLE 2:**
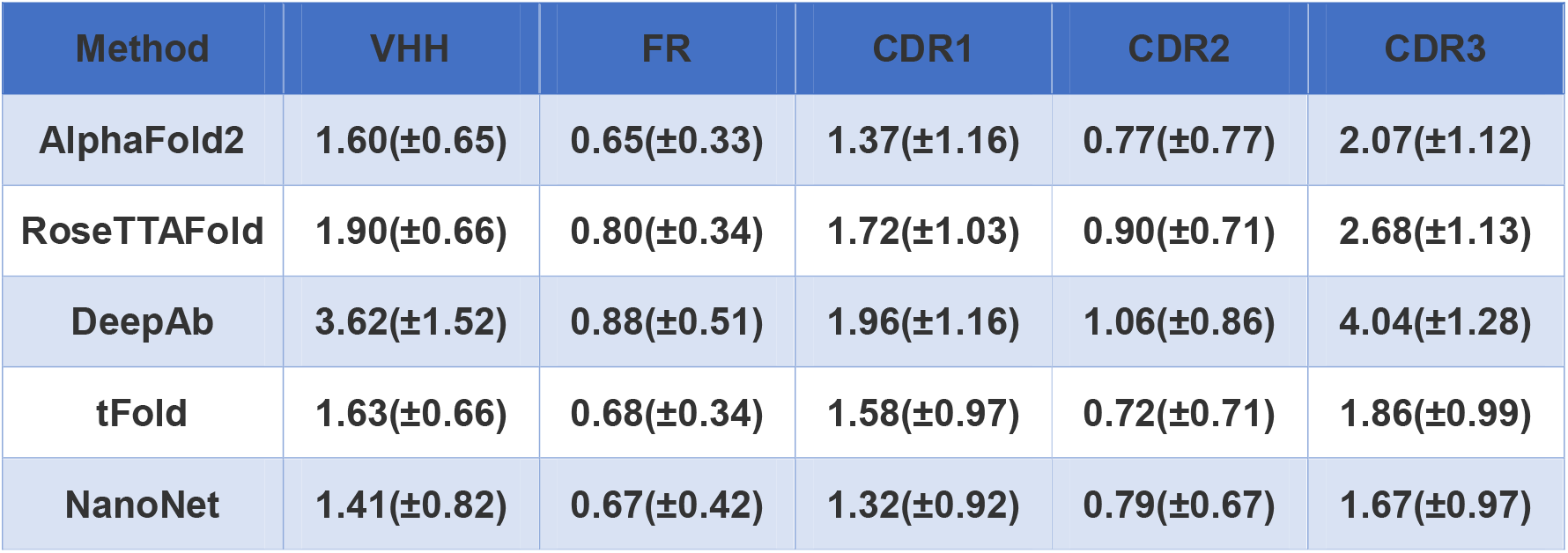
Mean RMSDs between the experimental and Cα-only predicted whole VHH and FR, CDR1, CDR2, and CDR3 structures for each method.

| Method | VHH | FR | CDR1 | CDR2 | CDR3 |
| --- | --- | --- | --- | --- | --- |
| AlphaFold2 | 1.60(±0.65) | 0.65(±0.33) | 1.37(±1.16) | 0.77(±0.77) | 2.07(±1.12) |
| RoseTTAFold | 1.90(±0.66) | 0.80(±0.34) | 1.72(±1.03) | 0.90(±0.71) | 2.68(±1.13) |
| DeepAb | 3.62(±1.52) | 0.88(±0.51) | 1.96(±1.16) | 1.06(±0.86) | 4.04(±1.28) |
| tFold | 1.63(±0.66) | 0.68(±0.34) | 1.58(±0.97) | 0.72(±0.71) | 1.86(±0.99) |
| NanoNet | 1.41(±0.82) | 0.67(±0.42) | 1.32(±0.92) | 0.79(±0.67) | 1.67(±0.97) |

The best performing method in the CDR2 ring region (Fig. 3A) was tFold with a mean RMSB of 0.72±0.71 Å, followed by NanoNet with a mean RMSD of 0.79±0.67 Å. DeepAb performed worst with a mean RMSD of 1.06±0.86 Å, followed by RoseTTAFold with a mean RMSD of 0.90±0.71 Å. AF2 was the middle performer with a mean RMSD of 0.77±0.77 Å.

NanoNet performed the best in the CDR3 ring region (Fig 3A) with a mean RMSD of 1.67±0.97 Å. However, NanoNet also had an outlier with a large RMSD in the CDR3 ring (Fig. A). The second best performing method was tFold with a mean RMSD of 1.86±0.99 Å. The worst performing method was still DeepAb, with a mean RMSD of 4.04±1.28 Å, followed by RoseTTAFold with a mean RMSD of 2.68±1.13 Å. AF2 was the middle-performing method with a mean RMSD of 2.07±1.12 Å (Table 2).

Comparing RMSDs with CDR3 loop length (Fig. 3B) showed that NanoNet performed best. Comparing CDR3 loop RMSDs among the different methods (Fig. 3C) also showed that NanoNet performed best. However, differences between NanoNet, tFold, and AF2 are small, with most points concentrated in the diagonal in these comparisons.

DeepAb always performed worse than the other methods. AF2 performed slightly better than RoseTTAFold. However, the differences between AF2 and tFold were small. NanoNet also performed slightly better than AF2. In addition, tFold performed better than RoseTTAFold. Moreover, NanoNet performed better than RoseTTAFold. Finally, NanoNet performed slightly better than tFold.

The CDR3 loop region is highlighted in the PyMOL structure visualization (Fig. 3D). AF2, RoseTTAFold, tFold, and NanoNet predictions were all relatively close to the experimental structure. However, the DeepAb prediction was far from the experimental structure for the CDR3 loop region.

When we compared the all-atom structures, tFold had the best TM scores followed closely by AF2. DeepAb performed worst with RoseTTAFold in the middle (Fig. 4A). This result is consistent with the RMSD comparisons. When we compare the Cα-only structures (Fig. 4B), NanoNet outperformed the other methods but had low-scoring outliers (Supplementary Table 3).

**FIGURE 4.**
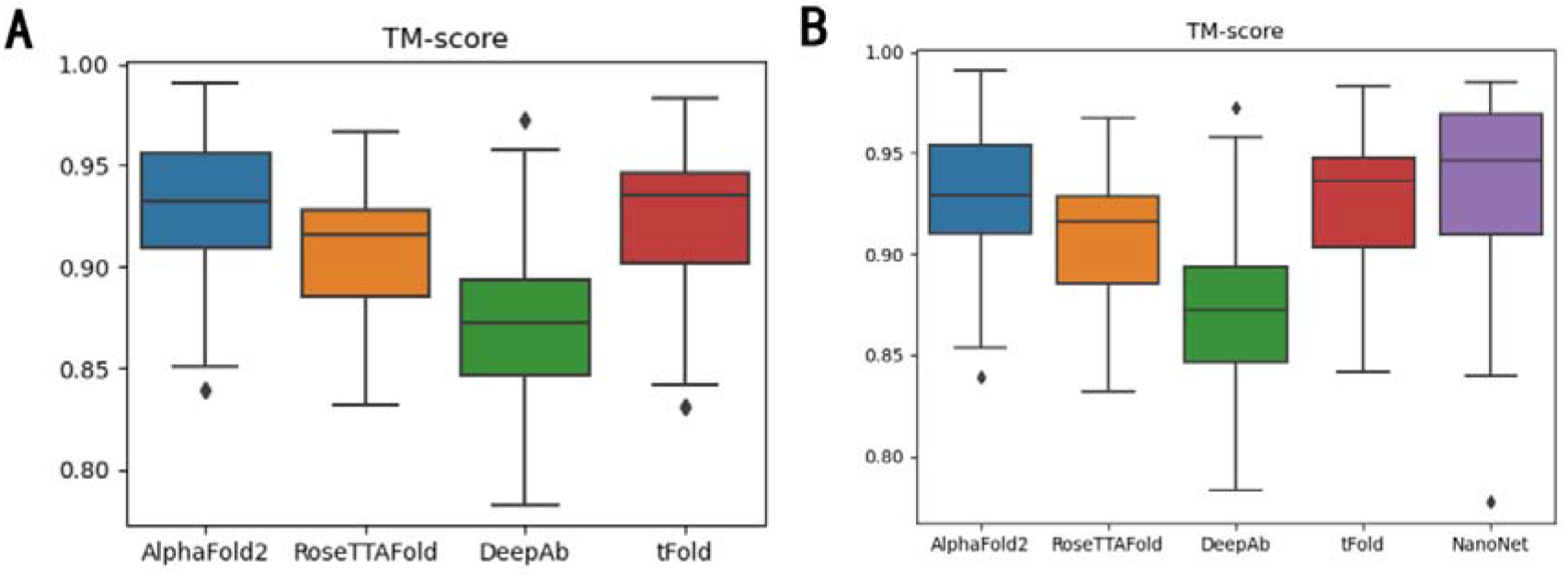
Boxplot comparison of TM scores for all-atom and Cα-only structures for each method. (A) Boxplots of TM scores for all-atom structures predicted by each method. (B) Boxplots of TM scores for the Cα-only structures predicted by each method.

Moreover, visualization of the predicted structures showed that RoseTTAFold performed poorly with Nbs whose CDR3 ring contained a β-turn or a similar structure (Supplementary Figure 3A), potentially explaining its poorer results compared to AF2.

### Uncertainty in DeepAb’s prediction results

We used DeepAb to make three predictions based on the amino acid sequence of a single H chain Nb and compared them. We found a degree of randomness in DeepAb’s predictions compared with the other methods, which all show good reproducibility (Supplementary Figure 3B). This randomness was more evident in the CDR3 ring but sometimes observed in other regions. We do not know whether this randomness is introduced during the selection of the optimal solution by DeepAb or reflects a bug in DeepAb’s prediction code.

## Discussion

### Performance on predictive accuracy

Our benchmarking of the accuracy of five deep learning Nb-folding algorithms in predicting VHH structures showed that NanoNet performed best, followed by tFold, AF2, and RoseTTAFold, all performing much better than DeepAb based on RMSDs (Figs. 2A and 3A). The main prediction errors were all in the CDR loop regions, particularly CDR3 loops, which had the greatest impact on overall prediction results, followed by CDR1 loops.

Surprisingly, the AF2 algorithm did not perform best in this highly variable region (Figs. 2B and 3B). However, the DeepAb algorithm, designed for traditional Ab Fv structure prediction, performed worst. Nevertheless, the NanoNet algorithm best predicted the overall framework and the local variable regions (Fig. 3A-B and Table 2). Incorporating an Nb folding database or algorithms such as NanoNet might improve the performance of protein folding algorithms such as AF2 in predicting highly variable regions. Importantly, we obtained the same method ranking order with TM scores (Fig. 4A-B).

### Computational time

It should be noted that NanoNet’s algorithm had the fastest model prediction speed. The prediction of the 60 input sequences only took 4.65 seconds since NanoNet can process them in parallel. In contrast, the average prediction time for a single Nb sequence using DeepAb on a local server with four NVIDIA Tesla V100 SXM2 graphics processing units in single-chain mode was around 3 minutes. The average prediction time for a single Nb sequence using AF2’s Colab notebook was around 5 minutes (https://colab.research.google.com/github/sokrypton/ColabFold/blob/main/AF2.ipynb). The average prediction time for a single Nb sequence using RoseTTAFold’s online Robetta platform (https://robetta.bakerlab.org, last accessed: July 14,2022) was 1—55 minutes, and each account is limited to 20 predictions per day.

The prediction time for a single Nb sequence using the tFold platform https://drug.ai.tencent.com/console/en/tfold?type=predict, last accessed: July 28, 2022) was usually 20—30 minutes but sometimes up to 9—16 hours. Inconveniently, only 10 tasks could be submitted per account per day, and only one result could be downloaded every 24 hours, greatly disadvantaging this online platform. We hope that more parallel submissions and downloads each day will be permitted in the future (Supplementary Figure 4).

### Robustness

At the algorithm’s functional level, NanoNet only supports the spatial prediction of five atoms (nitrogen [N], Cα, carbon [C], oxygen [O], and beta-carbon [Cβ]) for each amino acid backbone structure of the Nb (VHH) formed by the VH region of the common Y-shaped lgG Ab. DeepAb supports the prediction of the VL and VH double chains of the AB’s Fv or only the single chain VH or VL. The other three folding algorithms (AF2, RoseTTAFold, and tFold) support the prediction of the entire atomic protein structure of the Ab.

We also attempted to use the non-Nb protein hemoglobin (PDB ID: 2WY4) as input for the five algorithms to test their robustness. NanoNet was almost unusable for this hemoglobin protein prediction (RMSD=14.08Å). However, DeepAb’s prediction was even more unreasonable (RMSD=35.14Å). Nevertheless, AF2, RoseTTAFold, and tFold all predicted its 3D structure correctly (RMSD <3 Å; Supplementary Figure 5).

Folding predictions with Nb 1I3U showed that DeepAb, RoseTTAFold, and tFold did not allow wildcard “X”characters in the amino acid sequence. This “X” raised a ‘ValueError’ and halted the program. Therefore, we deleted the invalid ‘X’ character in the FASTA sequence to solve this problem. In addition, in the character study of input amino acid sequences, we found that some of the Nb FASTA files downloaded from the PDB database had polyhistidine tags^43,44^ at their C-terminal end. If the files with these polyhistidine tags were used directly for folding structure prediction, the RMSD prediction error from NanoNet would be 2.87±1.97 Å, which is much higher than when the tag was deleted (RMSD=1.38±0.82 Å).

### Relevance of the predicted outcomes and algorithm differences

Correlating RMSDs is a simple but efficient approach for determining whether a close relationship exists between pairs of algorithms. Indeed, the code of modern machine learning methods, especially protein folding-related deep learning methods, is more or less black-boxed. Therefore, we showed that AF2 showed the highest correlation with tFold (*r*=0.65), followed by AF2 with NanoNet (*r*=0.59), RoseTTAFold with tFold (*r*=0.57), and AF2 with RoseTTAFold (*r*=0.48; Fig. 5).

**FIGURE 5.**
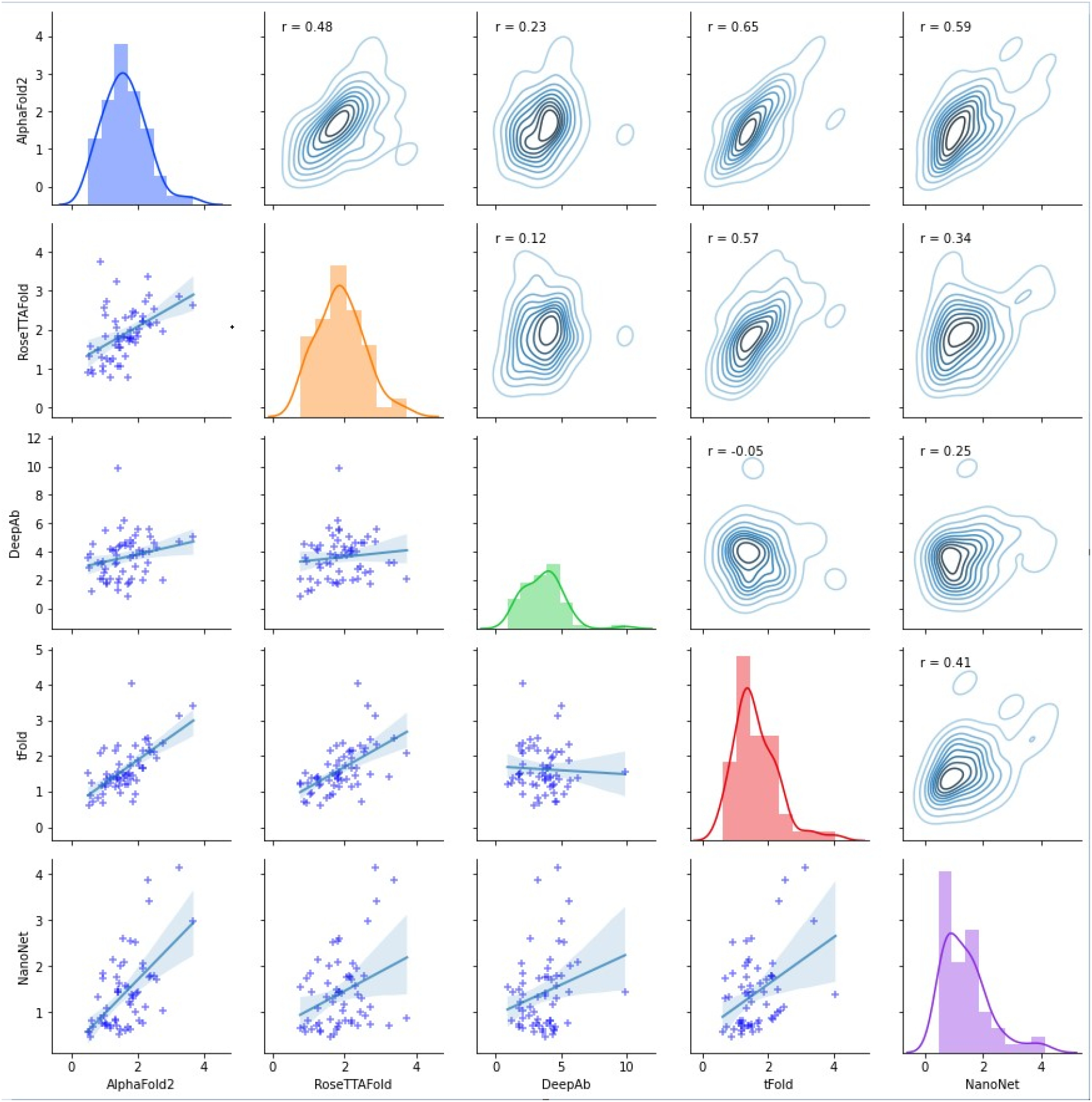
Pairwise Pearson’s correlations between Nb structure predictions by the five protein folding algorithms. Scatterplots below the matrix diagonal show the linear correlation of RMSDs between pairs of protein folding algorithms. Scatterplots above the matrix diagonal show the data’s kernel density between pairs of protein folding algorithms with the corresponding Pearson’s correlation (*r*). The kernel density of two algorithms with higher correlations is more concentrated along the diagonal, as shown by the nuclear density map of AF2 and tFold (top row, second from right; *r*=0.65). The RMSD density barplot for each algorithm is shown on the diagonal.

## Conclusions

AF2 performed best on FR in both all-atom and Cα-only structure predictions. In addition, AF2’s predictions for CDR1 and CDR2 were still in the top set. However, AF2’s predictions for CDR3 were only intermediate among methods. Overall, AF2 can accurately predict the overall structure of all-atom proteins, especially in stable regions such as Nb FRs.

RoseTTAFold and AF2 are both improvements on the AF1 algorithm.^17^ Their approaches are partially similar based on their reported algorithmic frameworks. However, their results were not well correlated. We expect that RoseTTAFold’s inclusion of a 3D track in the three-track block and differences in database selection underlie these differences.

The best all-atom Nb folding results were not obtained with these algorithms. Unexpectedly, the tFold algorithm developed by Tencent AI lab, released before RoseTTAFold and AF2, provided better results in the all-atom structure comparison. Like AF2, its folding step is divided into three homology modeling steps.^16^ The correlation values show that its predictions are highly similar to AF2 and RoseTTAFold. In addition, its prediction accuracies for CDR3s were significantly better than AF2 and RoseTTAFold, suggesting that tFold might be optimized explicitly for highly variable regions such as CDR3, consistent with Tencent AI Lab’s Ab drug discovery and development aims.

While NanoNet is a lightweight folding algorithm based on ResNet,^30^ its predictions were very similar to AF2, which used more sophisticated structures with an attention algorithm.^16,25^ Table 2 shows that it had the highest accuracy, especially in CDR3 regions, proving its ability in the Nb folding structure prediction and highlighting the importance of choosing a suitable training dataset for that specific sub-field. In addition, the lightweight NanoNet algorithm achieved remarkable speed due to its calculation based on the simplified Cα-only structure.

The DeepAb folding algorithm performed the worst in this study. DeepAb and its progenitor framework, DeepH3, are folding algorithms designed for traditional Ab Fv regions and are poorly suited for predicting Nb folding structures. Therefore, its poor performance is consistent with it being developed mainly for double-stranded Abs and its training datasets comprising mainly double-chained Abs. While DeepAb has a single-chain mode, it performed poorly in NB predictions. These results show how the algorithm and its training dataset can greatly influence prediction accuracy.

### Perspectives

New algorithms will continue to emerge in this newly emerging field with better architectures, prediction speed, and RMSD-based accuracy, which will greatly benefit protein and Nb 3D structure predictions, especially for high-variability such as CDR3.

## Materials and Methods

### Obtaining the PDB experimental file

We obtained the amino acid sequences and experimental PDB files of Nbs, removing duplicate entries and retaining those with the best resolution, from the official PDB website.^28^ Next, we inputted these Nb amino acid sequences into the five structure prediction algorithms to generate prediction PDB files. Then, the experimental PDB files were used as the gold standard against which predicted PDB files were compared to calculate RMSDs and TM scores.

### RMSB calculation

We uploaded the experimental and prediction PDB files into PyMOL (https://github.com/schrodinger/pymol-open-source) to calculate RMSDs for each prediction algorithm. PyMOL automatically aligns the atoms between the predicted and experimental PDB files. When calculating RMSDs with the default outlier rejection cutoff of 2.0 Å in PyMOL’s align parameter, atoms in the two input PDB files with large distance differences are removed before the RMSD calculation. This default filtering threshold removes outliers automatically but can lead to indistinguishable RMSD results due to removing all abnormal values. To ensure fair and reasonable results and to retain the maximum number of atoms in the experimental PDB file for the RMSD calculation, we set this cutoff to 10. This threshold ensured that atoms with differences up to 10 were not removed and that RMSDs were more accurate and reliable.

### CDR rings segmentation by the international immunogenetics information system (IMGT) scheme

The Nb protein sequence was divided into seven regions (FR1, CDR1, FR2, CDR2, FR3, CDR3, and FR4) according to the IMGT annotation method.^45^

### PDB file partitioning based on CDR loops

The IMGT annotation method divided Nb amino acid sequences into seven different FR (FR1–4) and CDR (CDR1–3) regions. Based on the information on the amino acid sequences of these seven regions, we also segmented the experimental PDB files obtained from the official PDB website into the same seven regions. We also segmented the processed experimental structure PDB files containing all and Cα-only atoms into CDR regions. After segmentation, each FR or CDR loop region was a separate PDB file. That is, Nb PDB files were segmented into seven PDB files, one for each FR or CDR. Since the NanoNet prediction files only contained 3D coordinates of Cα atoms,^35^ we filtered the PDB files for the other four methods, retaining only Cα atoms and information on the five atoms (N, Cα, C, O, and Cβ) to enable comparisons with NanoNet.

### RMSD calculation for the CDR loops

PyMOL enabled us to align all atoms and residues and to calculate RMSDs, which helped us automatically compare parts of the predicted and experimental structures. We directly calculated RMSDs, comparing the predicted PDB files generated by each algorithm to the experimental structure PDB file for each segmented CDR region. This approach is comparable to CDR partitioning the predicted PDB files followed by RMSD calculation with the partitioned experimental PDB files.

### TM score calculation

The TM score was calculated using TM-align.^41^ The experimental and predicted PDB files were passed to the TM-align method to calculate the TM score.

### Comparison of NanoNet with other methods

While the NanoNet output contains only Cα coordinates, the output of the other algorithms also contains non-Cα coordinates. Therefore, for comparisons between NanoNet and other algorithms, we filtered their files, retaining only the Cα coordinates for RMSD calculation; only the information of five atoms (N, Cα, C, O, Cβ) was retained in the PDB file. Although this method deleted other residue atoms, we could still obtain a basic spatial structure of the predicted results. Only retaining the Cα atom had little effect on the Nb spatial structure. This transformation did not change the 3D coordinates of atoms in the PDB file. Only the PyMOL visualization results sometimes differed slightly from those based on all atoms (Supplementary Figure 6).

### Ramachandran plot

We used the Drug RamachanDraw program to draw Ramachandran plots (https://github.com/alxdrcirilo/RamachanDraw).^42^ The Ramachandran plot was drawn by passing the prediction PDB file to the Ramachandran plotting tool in Python.

## Supporting information

Ramachandran_plot

## Acknowledgments

We thank Chunhai Xue, Leixin Zhu, Zelong Li, and Huanqing Long for data collection and all the members of the Wang lab for helpful discussions.

## Authors’ contributions

XinW conceived and supervised the study. SL, ZQL, and ZW performed the computational analyses. SL and ZQL interpreted the data. SL, ZQL, ZW, and XinW wrote the paper. FH, XuW, ZT, RH, ZYL, YC, BH, and XinW revised the paper. All authors read and approved the final manuscript.

