## Supplementary Figures for "A benchmark study of protein folding algorithms on nanobodies"

**Supplementary Figure 1. Outlier (PDB ID: 3K3Q) in CDR1 by AF2.**

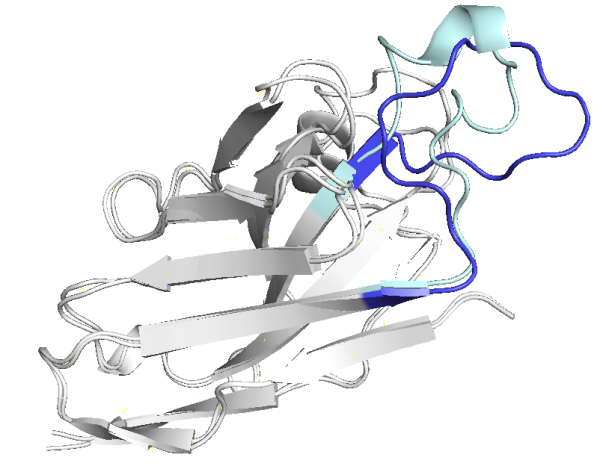

Blue: AF2 result for CDR1.

Cyan: Experimental results for CDR1.

Silver: Experimental results for other regions.

**Supplementary Figure 2. Boxplots of CDR lengths and their correlations with CDR RMSD, average CDR RMSD, and VHH RMSD.**

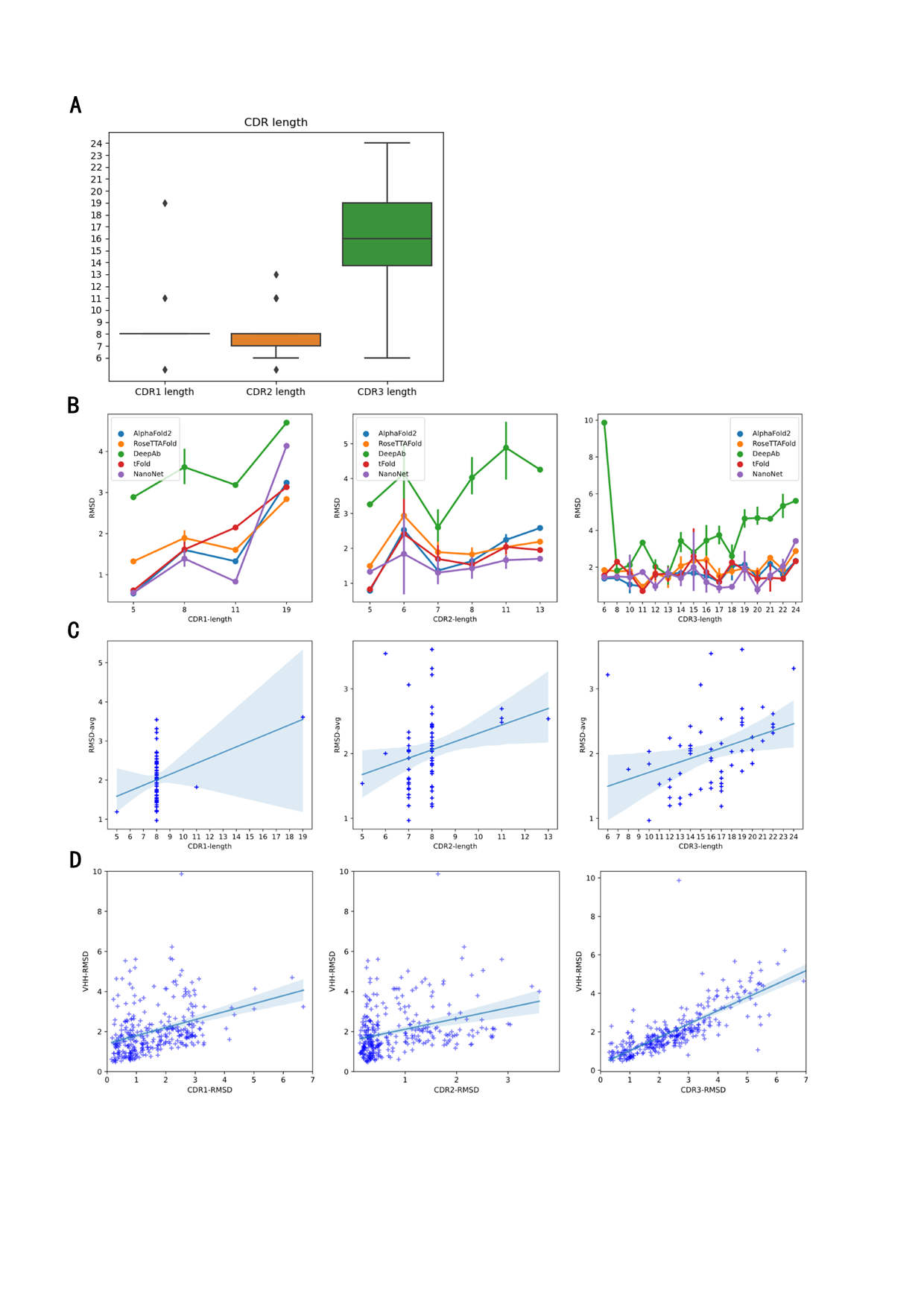

Figure A: Boxplot of CDR1, CDR2, and CDR3 lengths.

Figure B: Line graphs of each CDR’s length with the mean RMSD of each VHH predicted by the five algorithms.

Figure C: Scatterplots for each CDR’s length with average RMSD of each Nb predicted by the five algorithms.

Figure D: Scatterplots of the average RMSD of each CDR predicted by the five algorithms with the average RMSD of each VHH predicted by the five algorithms.

**Supplementary Figure 3. Visual comparison.**

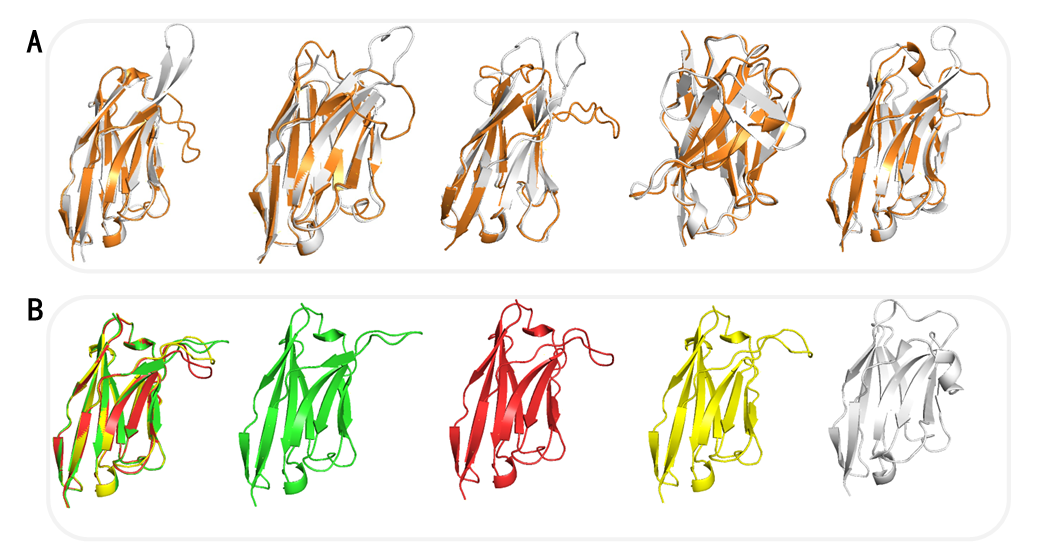

Figure A: Inaccuracy of RoseTTAFold predictions (orange) on β-turn structures (or similar structures) in the CDR3 loop (from left to right, PDB IDs: 4IDL, 4B50, 1KXV, 4HEP, and 5HVF) compared to the experimental structure (silver).

Figure B: Uncertainty of DeepAb’s three predictions for the same sequence (PDB ID: 4POY; the first attempt is shown in green, the second attempt in red, and the third attempt in yellow) compared to the experimental structure (silver).

**Supplementary Figure 4. Boxplot of computational time of each algorithm.**

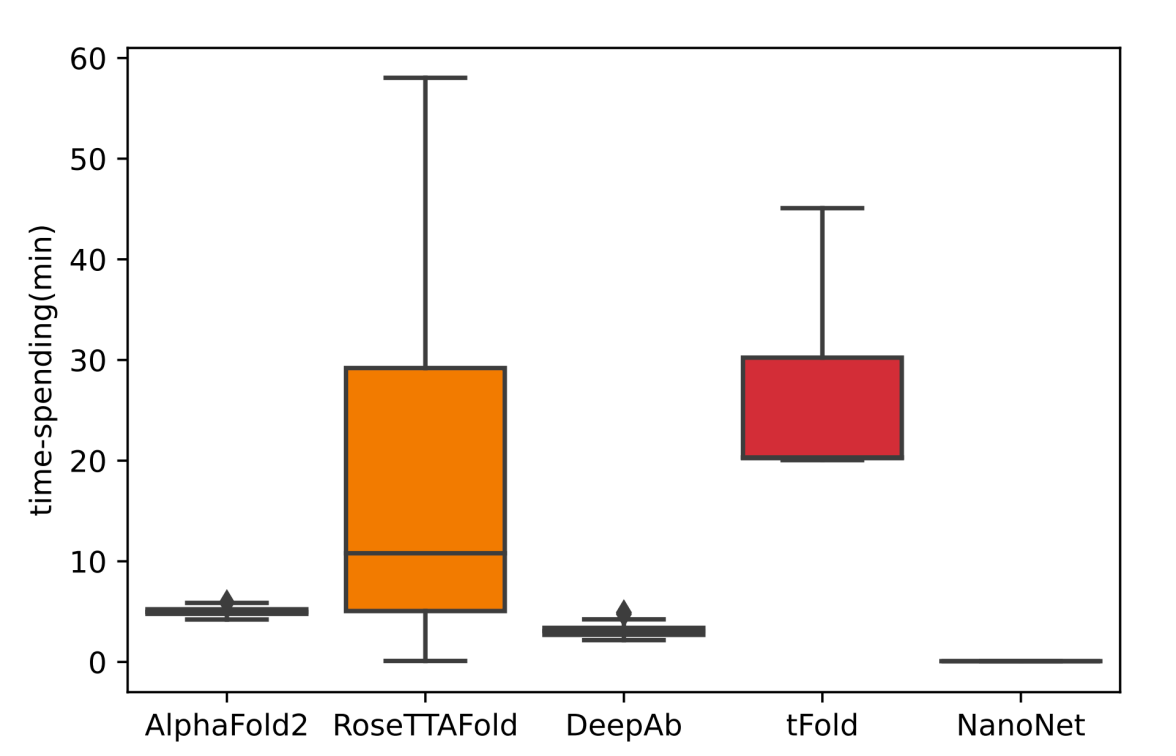

**Supplementary Figure 5. Hemoglobin (PDB ID: 2WY4) predictions by the five algorithms to evaluate their robustness.**

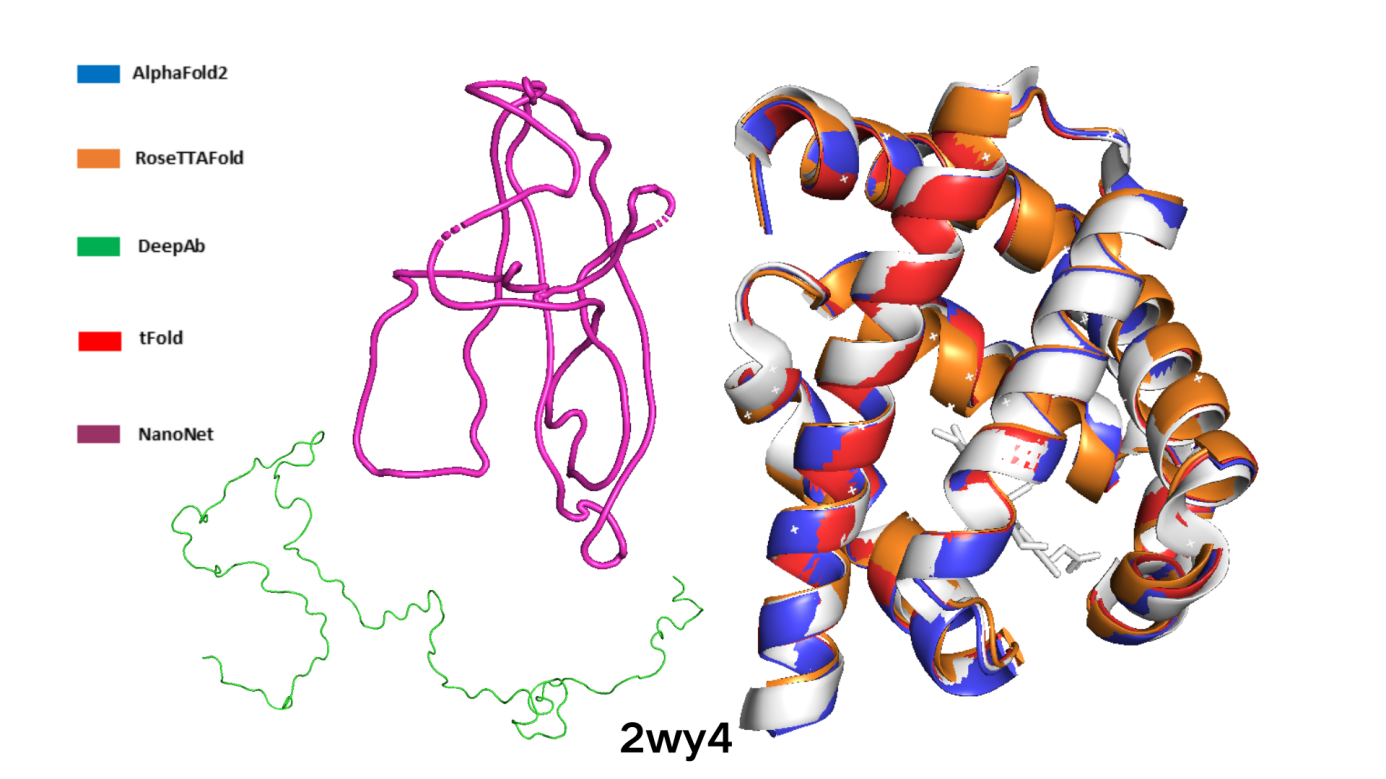

**Supplementary Figure 6. Visual comparison of the all-atom (PDB ID: 2VYR) and Cα-only (PDB ID: 4z9k) structures.**

**Supplementary Figure 6 Alt Text [37 words].** There are two visual comparisons**.** In the first, the two structures overlap completely (PDB ID: 2VYR). In the second, the all-atom structure (silver) differs slightly from the Cα-only structure in the light cyan area (PDB ID: 4Z9K).

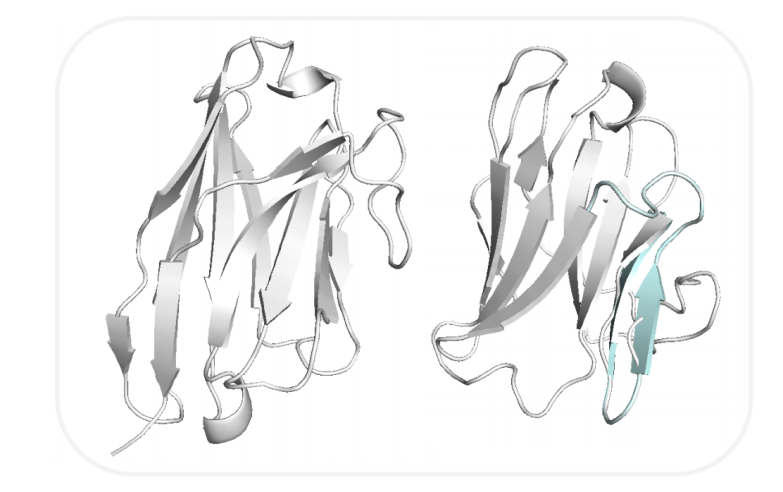
