## Supplementary figures and images for "A benchmark study of protein folding algorithms on nanobodies"

### 1i3v_plot_NanoNet.png

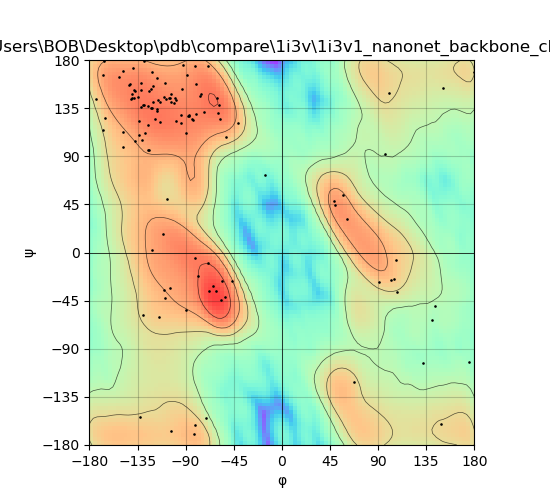

### 1i3v_plot_RoseTTAFold.png

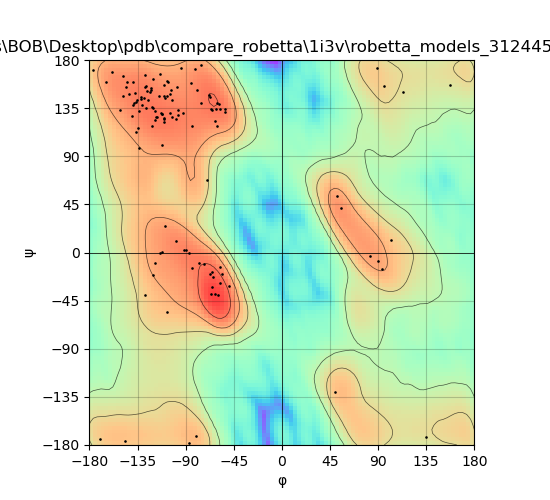

### 1kxv_plot_NanoNet.png

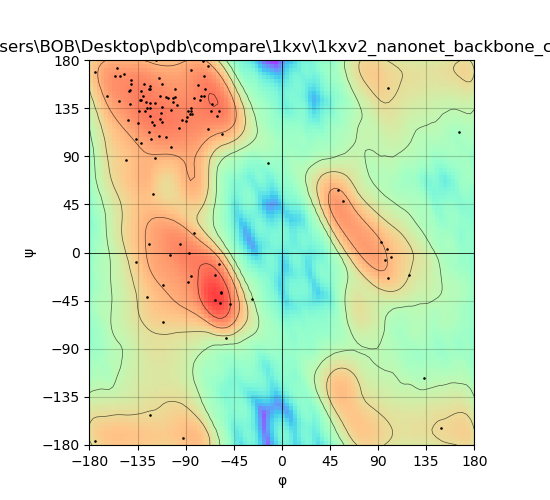

### 1kxv_plot_RoseTTAFold.png

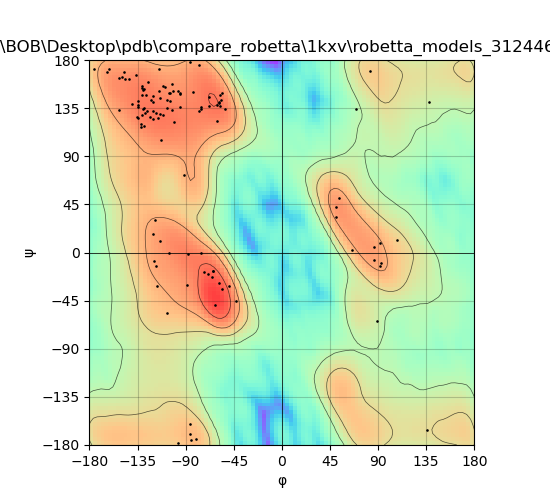

### 1op9_plot_NanoNet.png

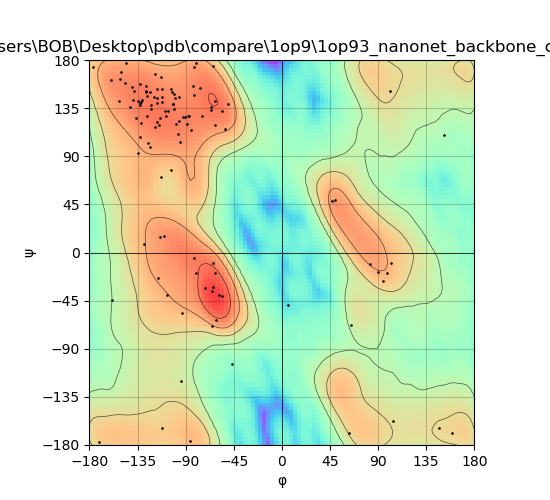

### 1op9_plot_RoseTTAFold.png

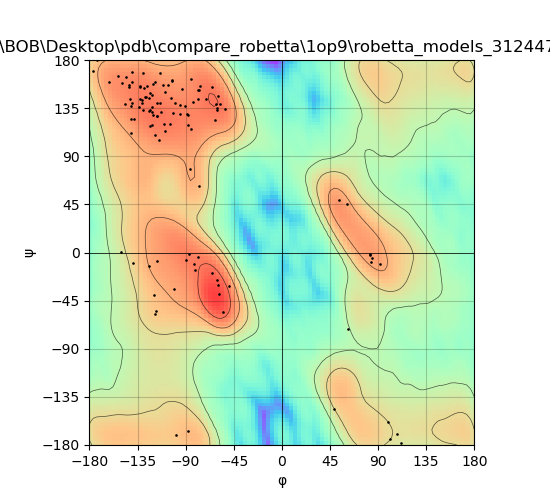

### 1qd0_plot_NanoNet.png

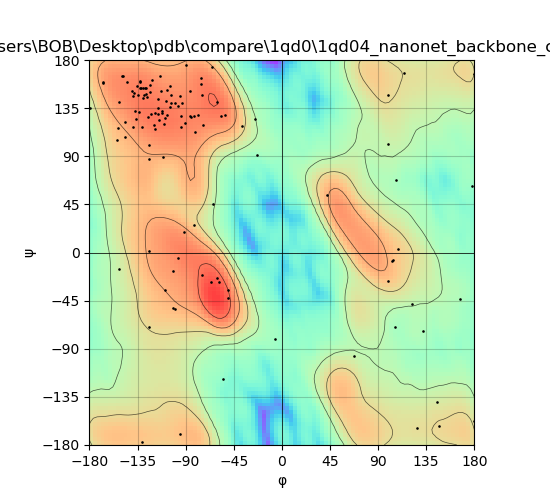

### 1qd0_plot_RoseTTAFold.png

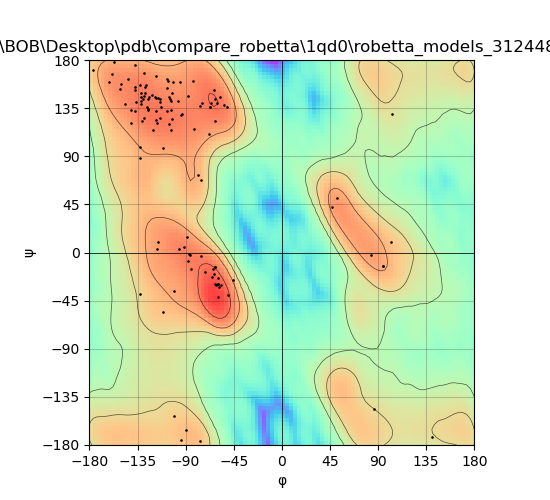

### 2vyr_plot_RoseTTAFold.png

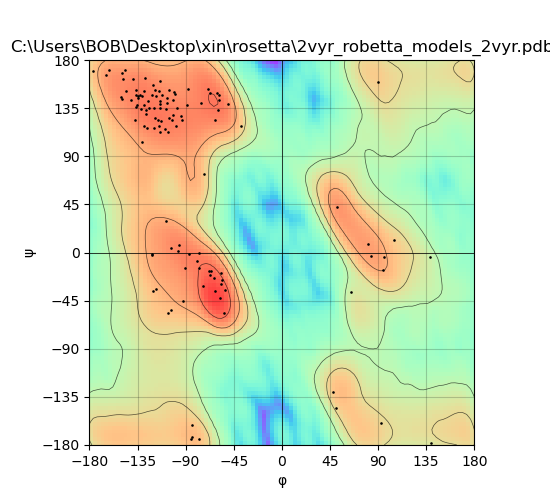

### 2vyr_plot_tFold.png

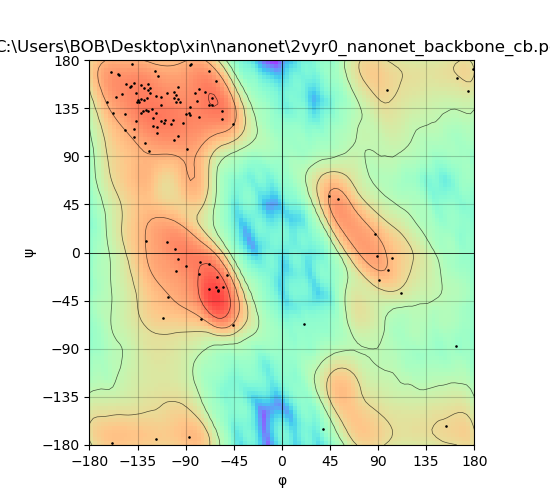

### 2xxc_plot_NanoNet.png

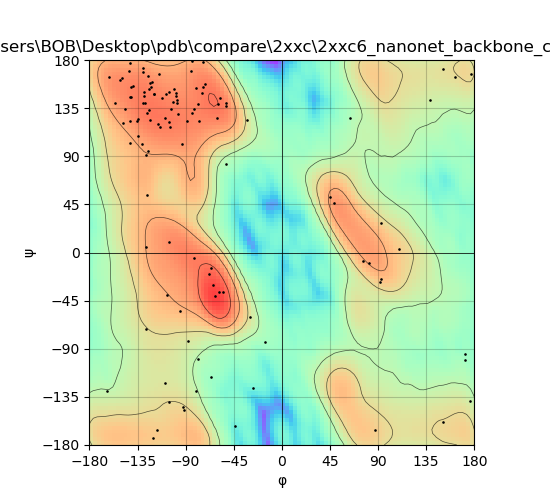

### 2xxc_plot_RoseTTAFold.png

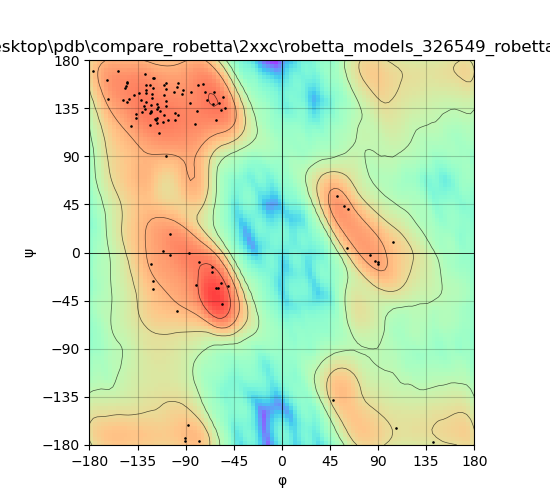

### 3jbc_plot_NanoNet.png

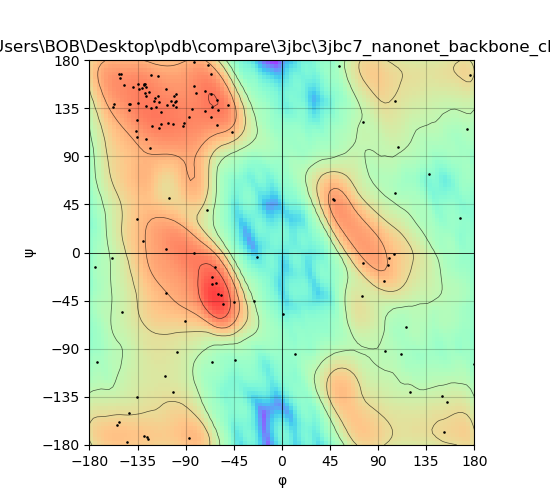

### 3jbc_plot_RoseTTAFold.png

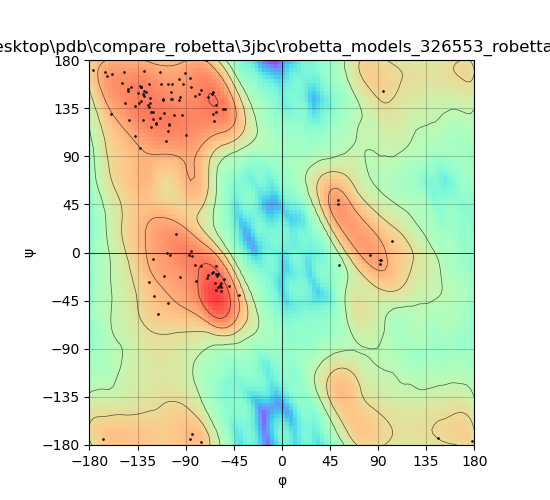

### 3jbd_plot_NanoNet.png

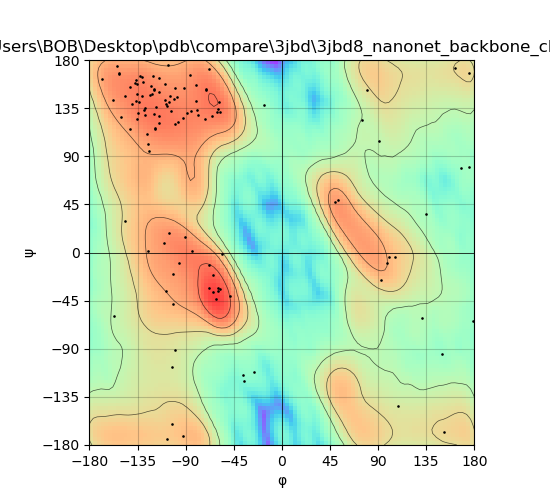

### 3jbd_plot_RoseTTAFold.png

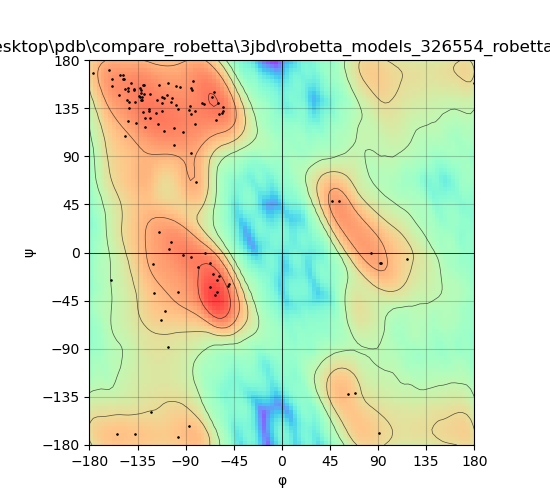

### 3k3q_plot_NanoNet.png

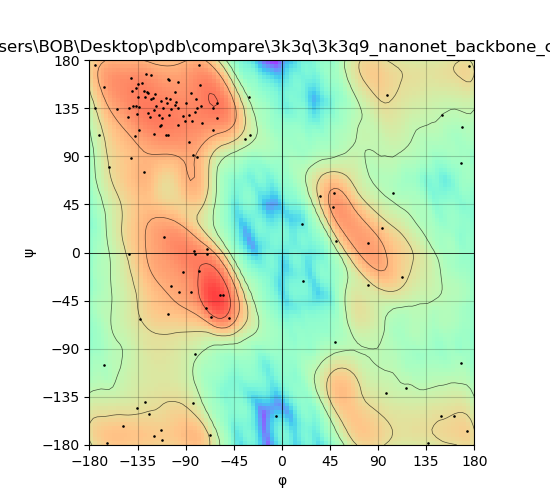

### 3k3q_plot_RoseTTAFold.png

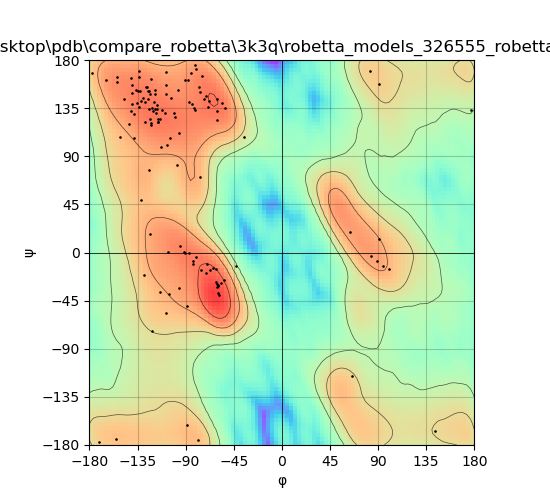

### 3k81_plot_NanoNet.png

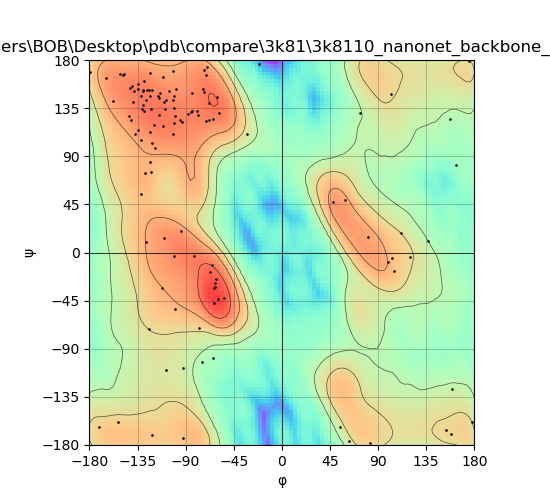

### 3k81_plot_RoseTTAFold.png

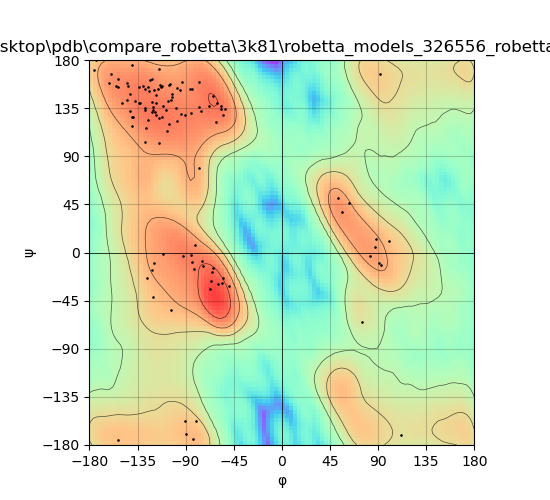

### 3qsk_plot_NanoNet.png

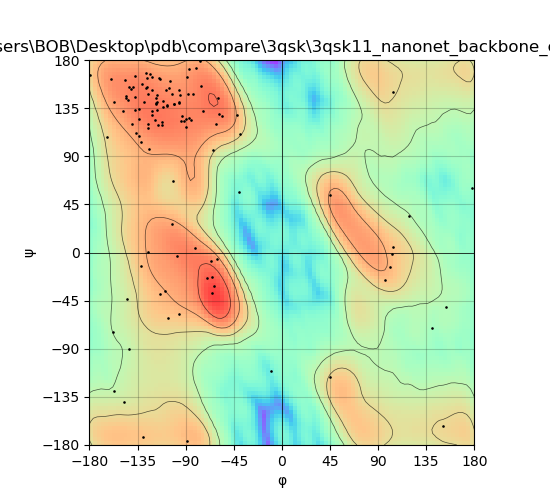

### 3qsk_plot_RoseTTAFold.png

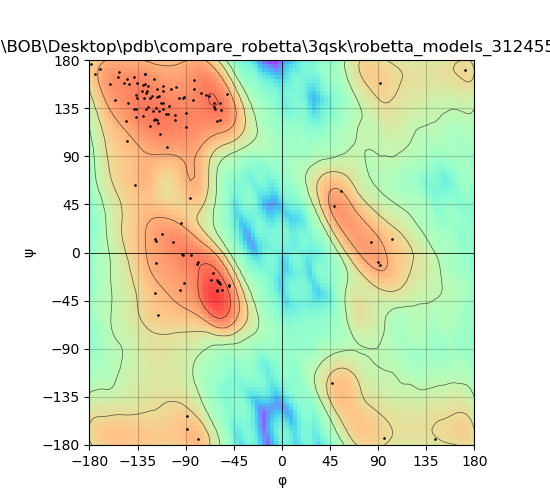

### 3qxt_plot_NanoNet.png

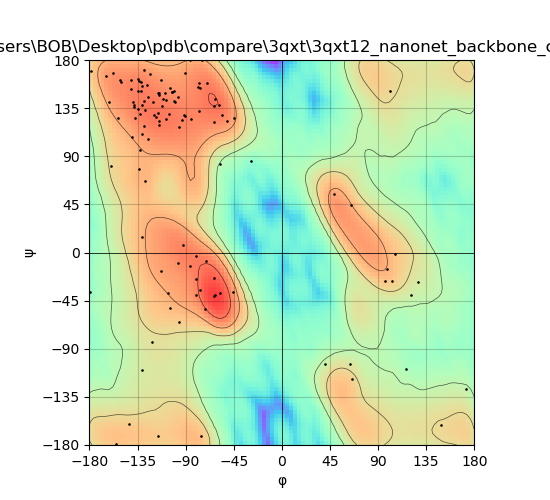

### 3qxt_plot_RoseTTAFold.png

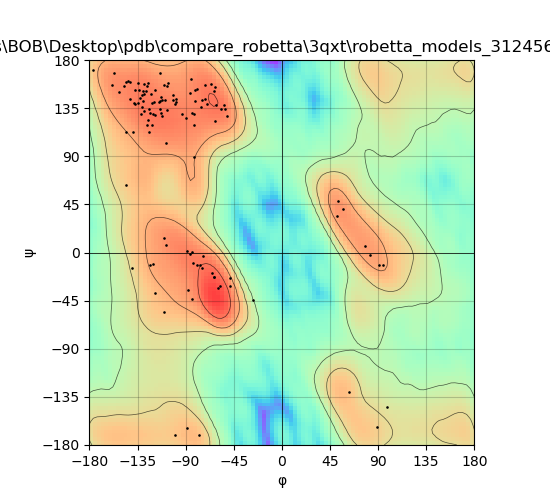

### 3r0m_plot_NanoNet.png

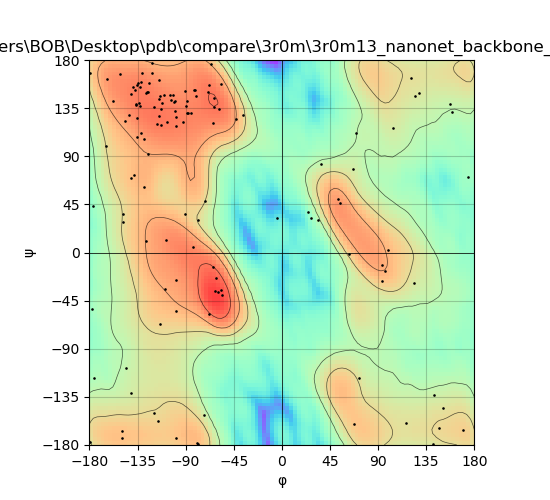

### 3r0m_plot_RoseTTAFold.png

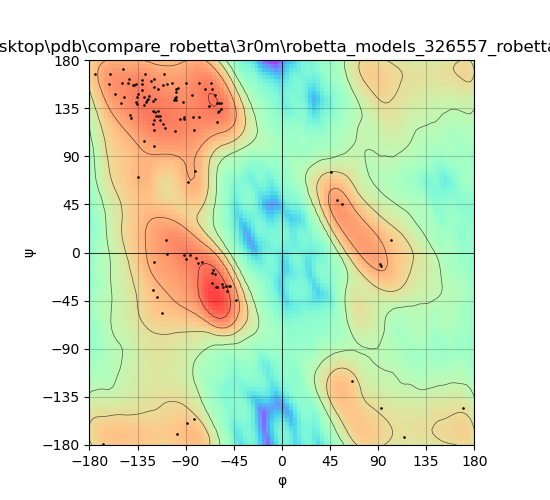

### 3sn6_plot_NanoNet.png

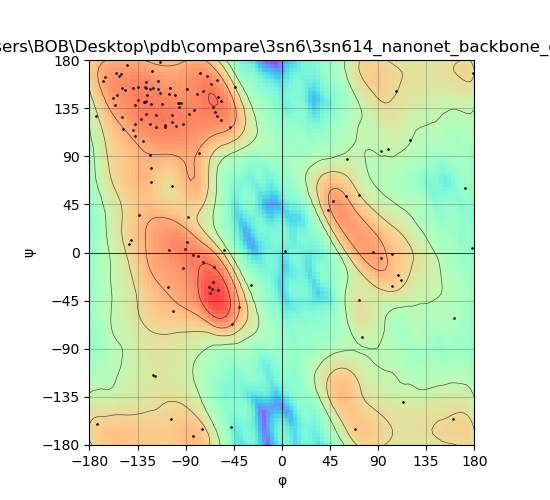

### 3sn6_plot_RoseTTAFold.png

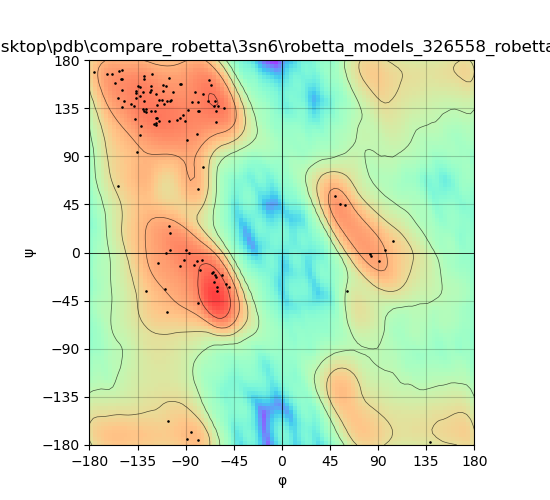

### 3stb_plot_NanoNet.png

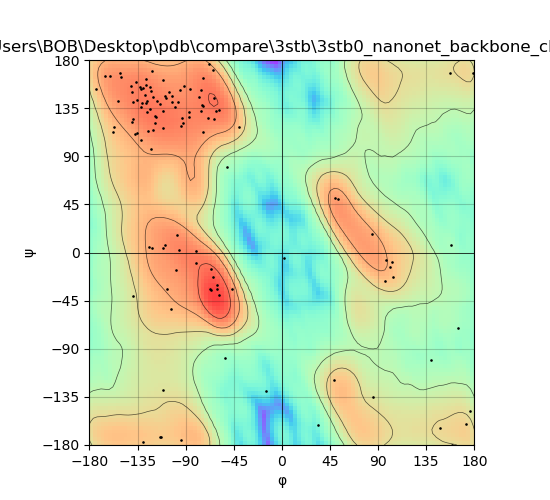

### 3stb_plot_RoseTTAFold.png

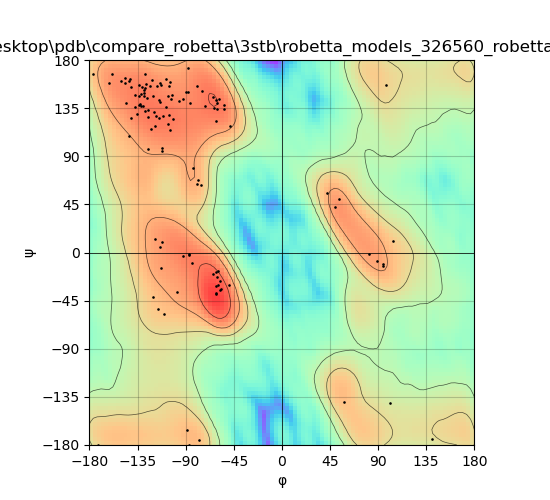

### 4b50_plot_NanoNet.png

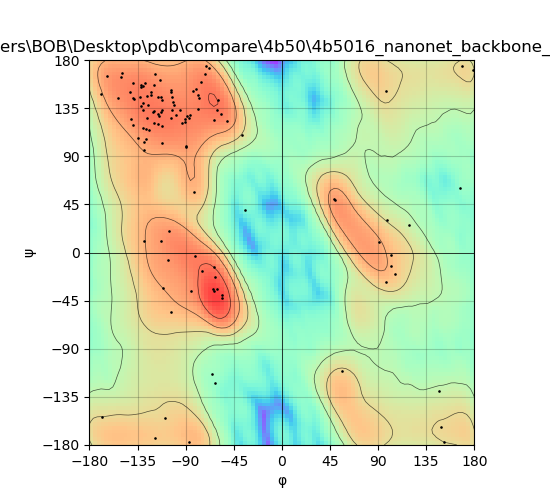

### 4b50_plot_RoseTTAFold.png

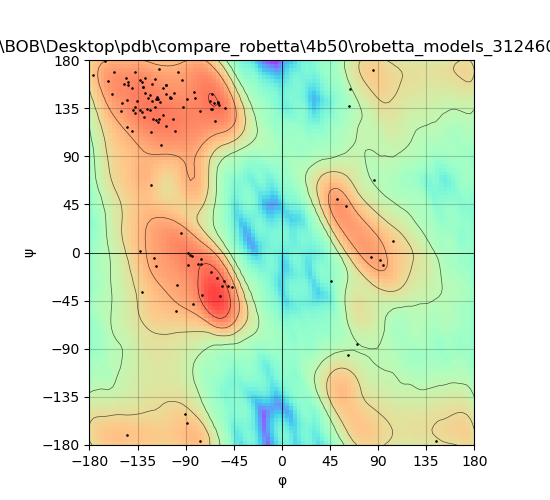

### 4dk3_plot_NanoNet.png

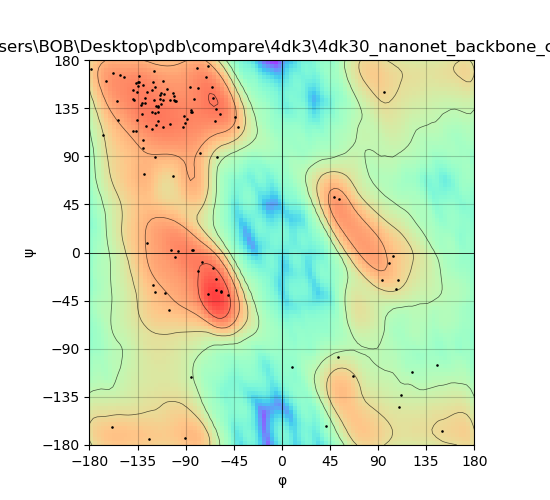

### 4dk3_plot_RoseTTAFold.png

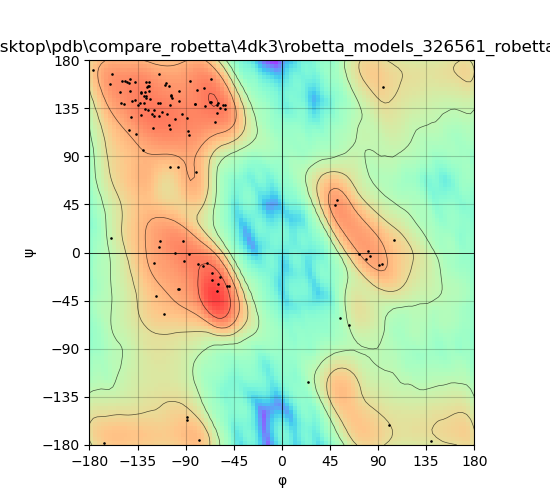

### 4hem_plot_NanoNet.png

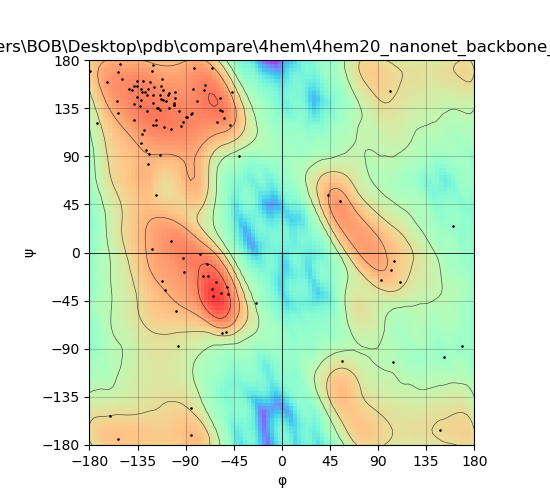

### 4hem_plot_RoseTTAFold.png

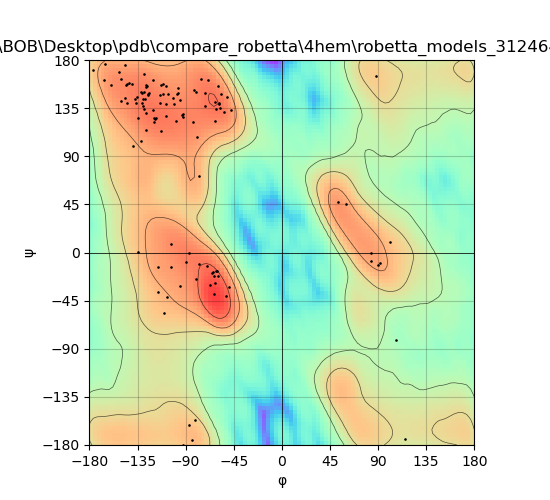

### 4hep_plot_NanoNet.png

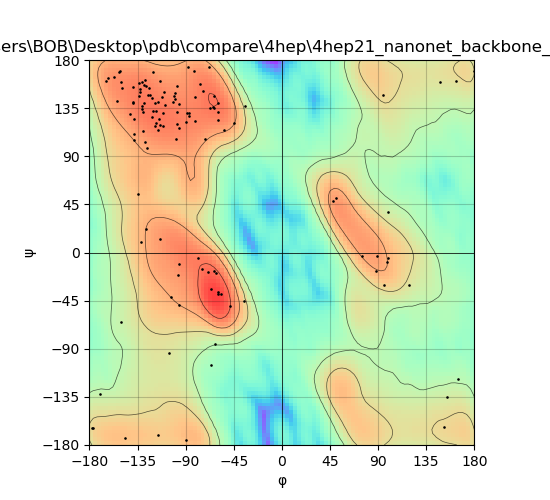

### 4hep_plot_RoseTTAFold.png

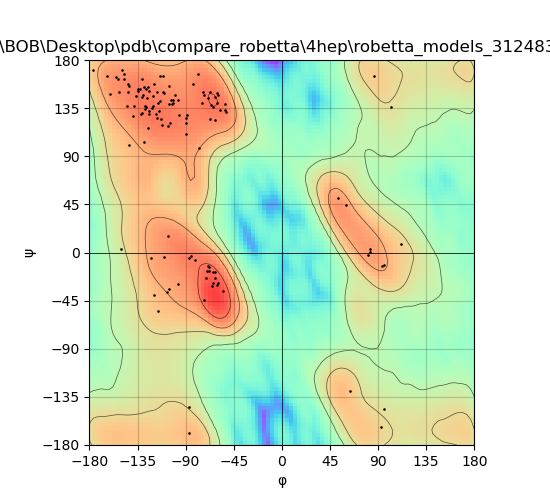

### 4idl_plot_NanoNet.png

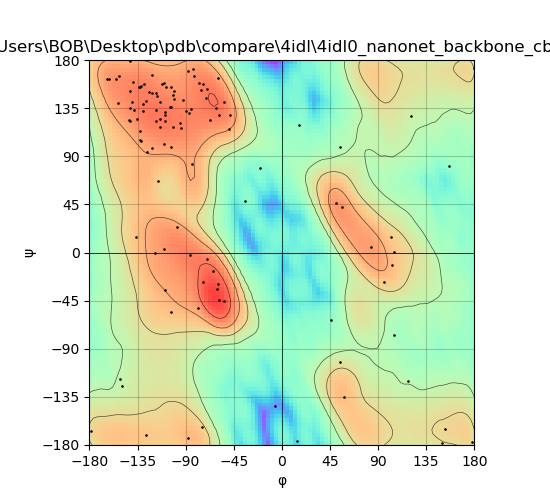

### 4idl_plot_RoseTTAFold.png

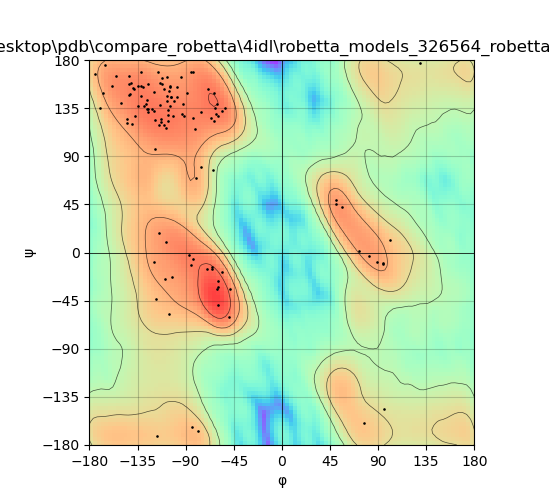

### 4lgp_plot_NanoNet.png

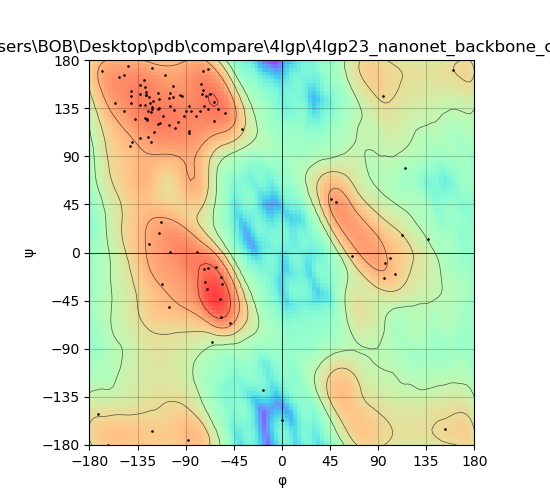

### 4nbx_plot_NanoNet.png

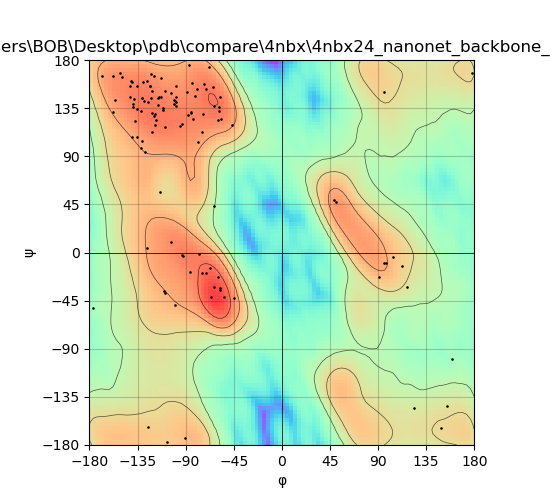

### 4nc0_plot_NanoNet.png

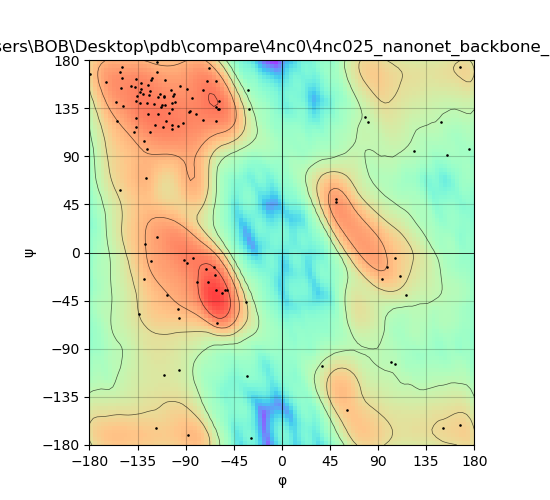

### 4nc2_plot_NanoNet.png

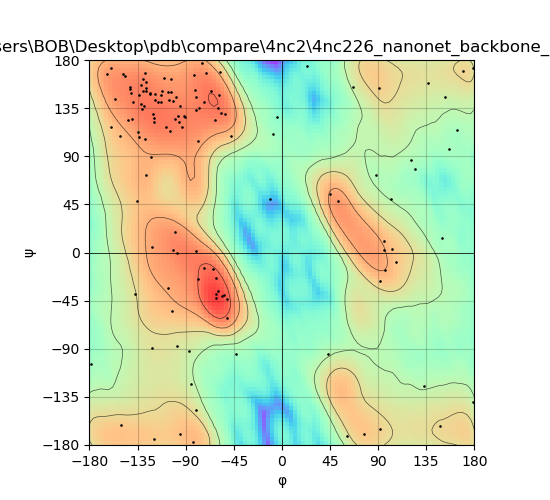

### 4orz_plot_NanoNet.png

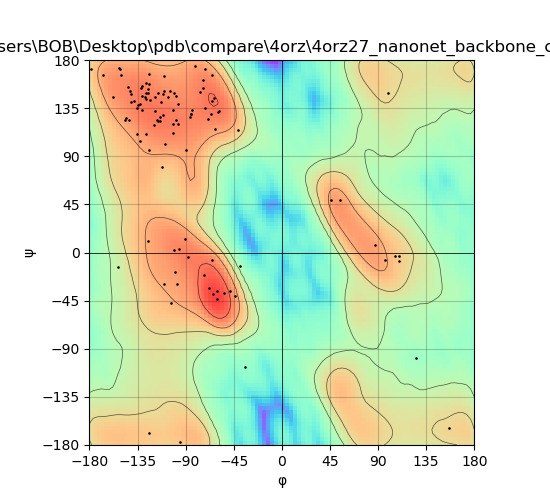

### 4pou_plot_NanoNet.png

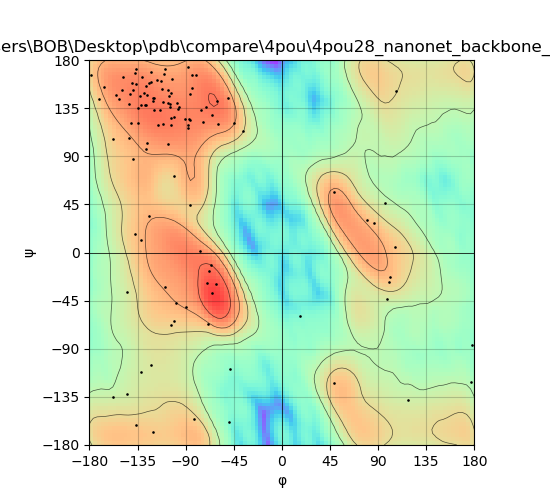

### 4poy_plot_NanoNet.png

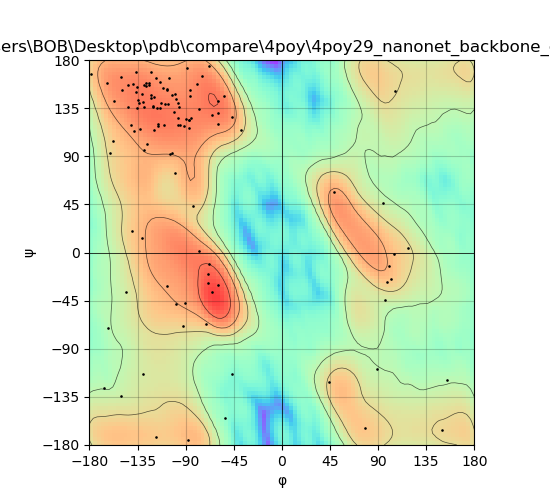

### 4ppt_plot_NanoNet.png

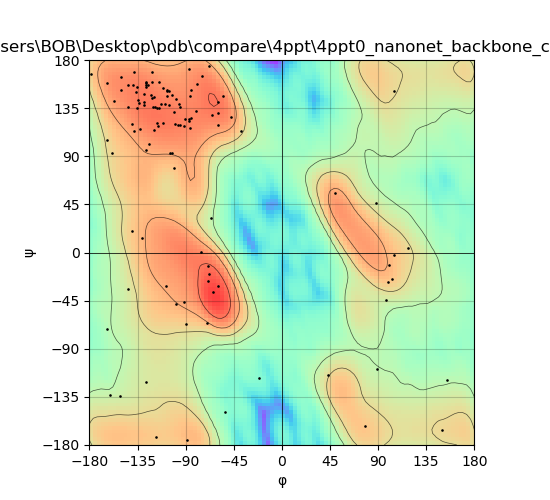

### 4tvs_plot_NanoNet.png

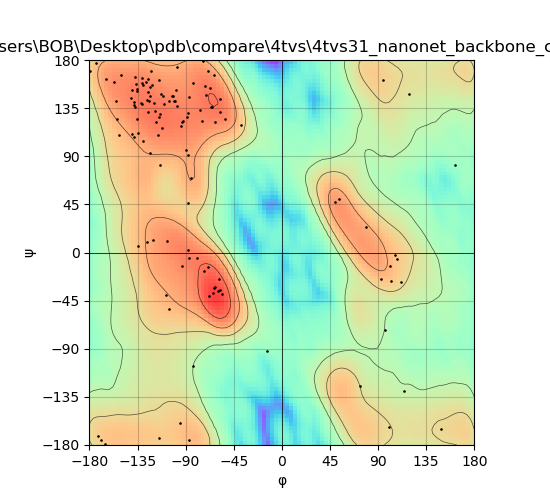

### 4tyu_plot_NanoNet.png

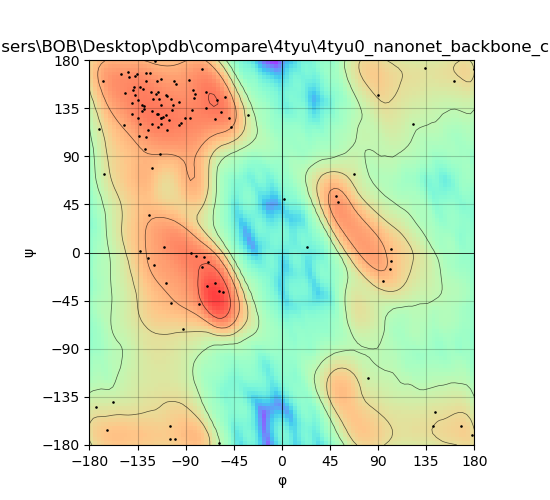

### 4u05_plot_NanoNet.png

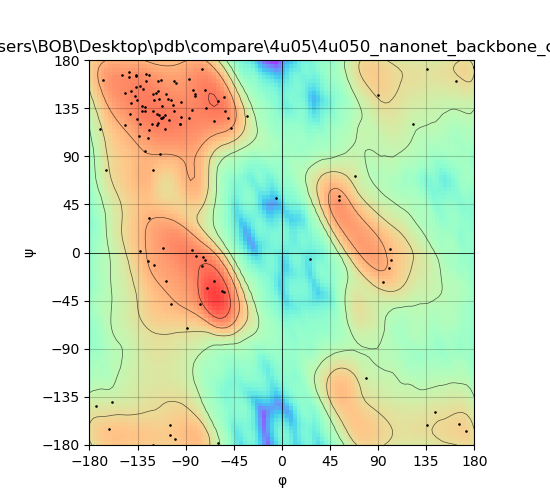

### 4w2q_plot_tFold.png

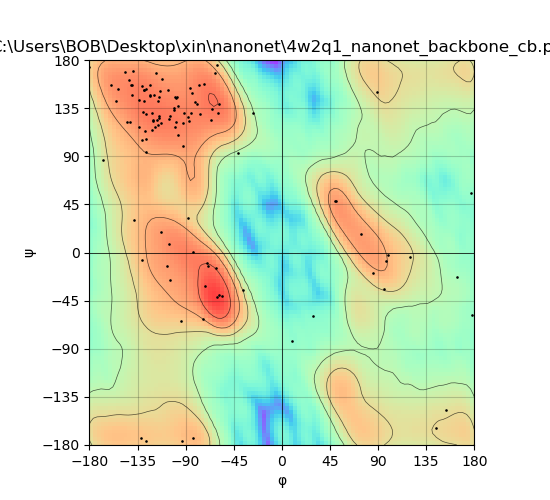

### 4w70_plot_NanoNet.png

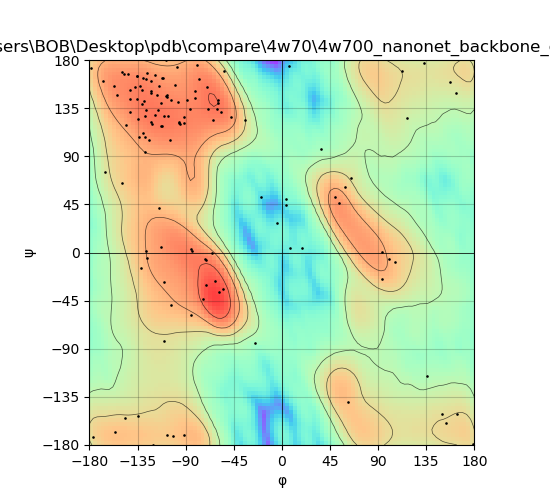

### 4wem_plot_NanoNet.png

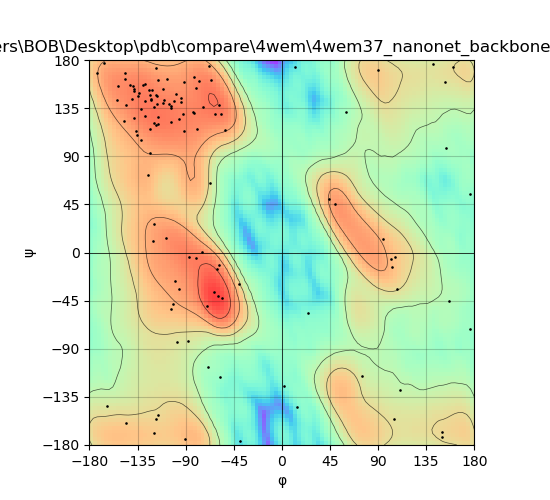

### 4yga_plot_tFold.png

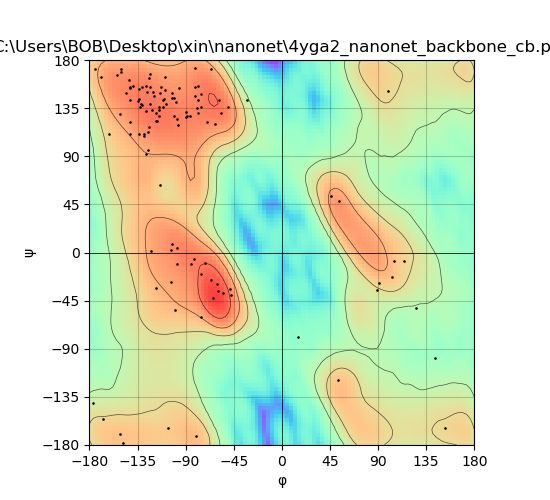

### 4z9k_plot_NanoNet.png

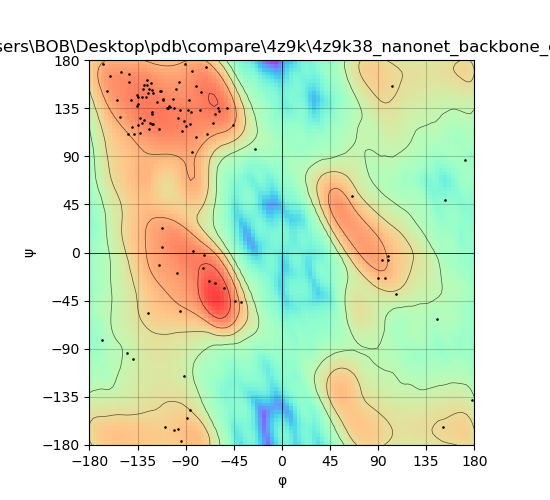

### 5boz_plot_NanoNet.png

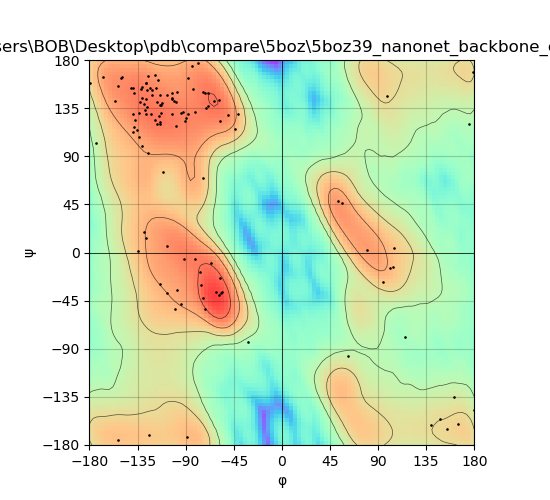

### 5e1h_plot_NanoNet.png

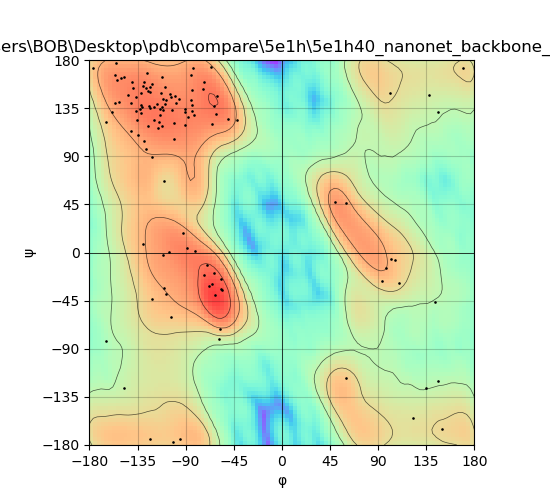

### 5hvf_plot_NanoNet.png

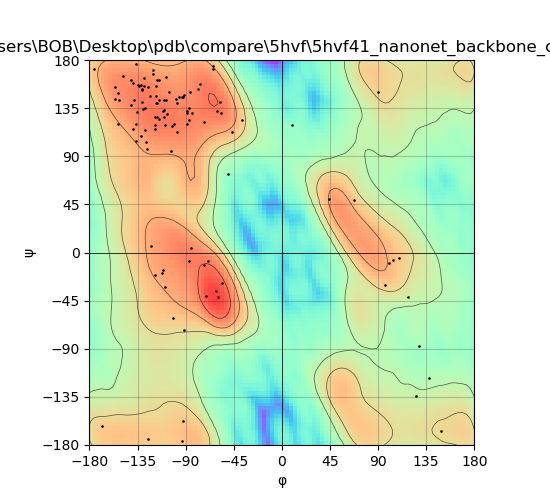

### 5j1s_plot_NanoNet.png

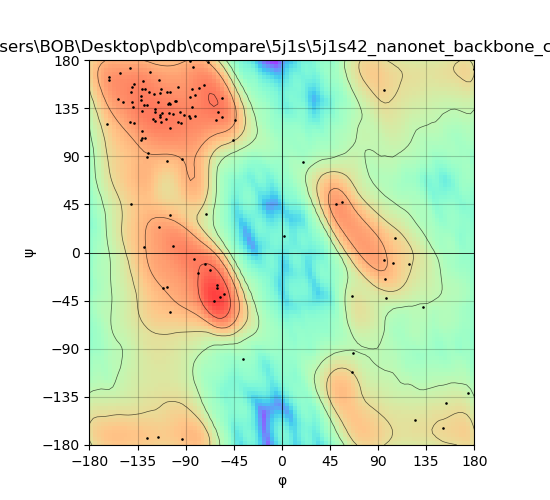

### 5j57_plot_DeepAb.png

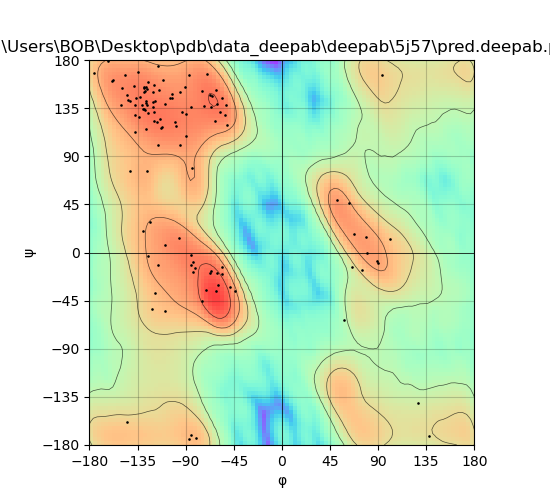

### 5j57_plot_NanoNet.png

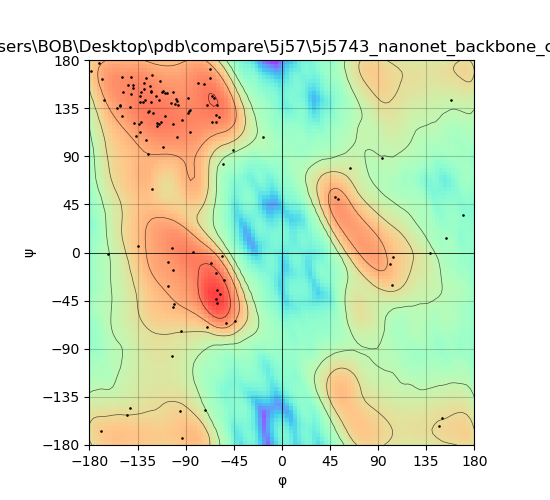

### 5jmo_plot_DeepAb.png

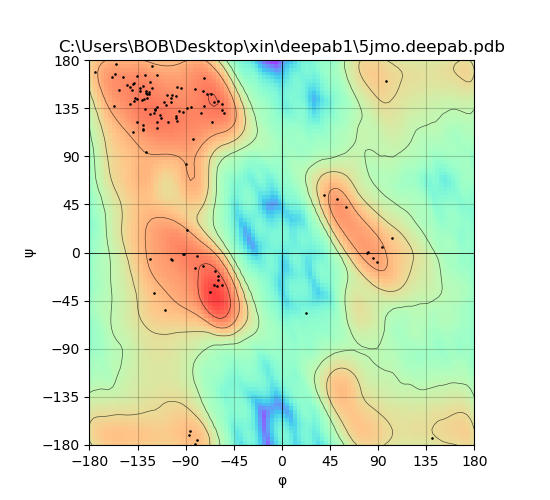

### 5jmo_plot_tFold.png

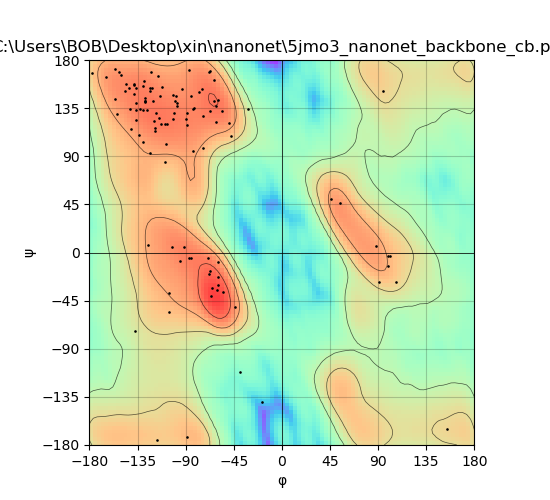

### 5nlu_plot_DeepAb.png

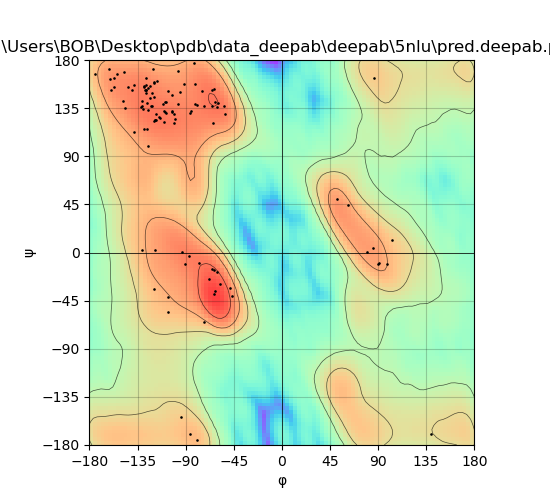

### 5nlu_plot_NanoNet.png

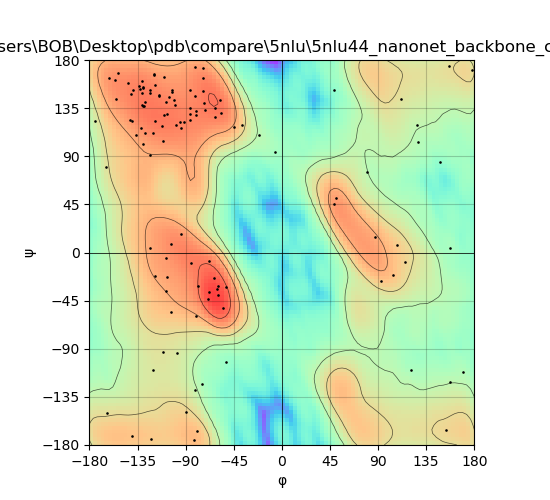

### 5sv3_plot_DeepAb.png

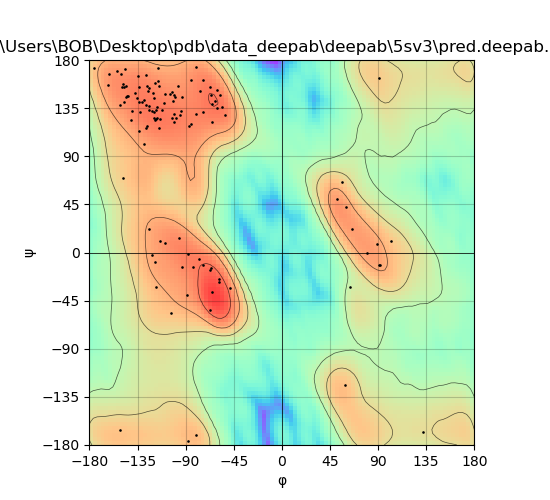

### 5sv3_plot_NanoNet.png

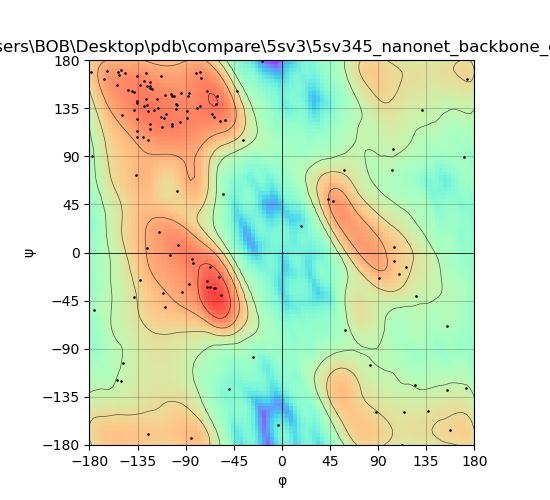

### 5vlv_plot_DeepAb.png

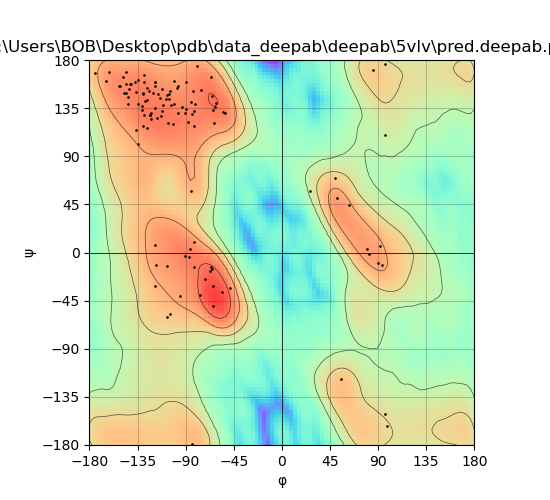

### 5vlv_plot_NanoNet.png

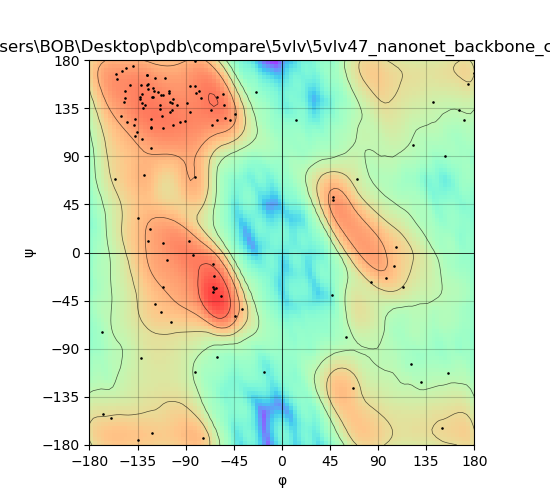

### 5vm6_plot_DeepAb.png

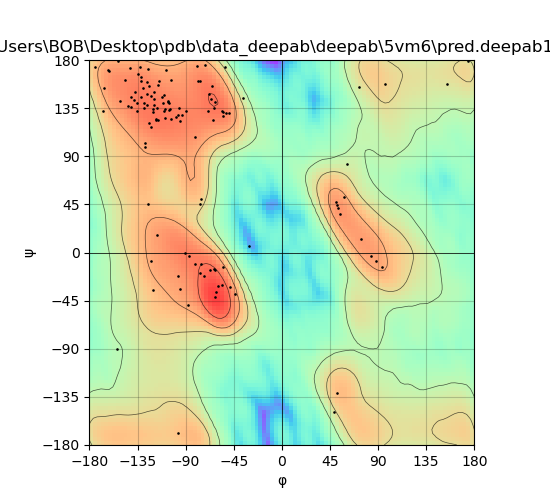

### 5vm6_plot_NanoNet.png

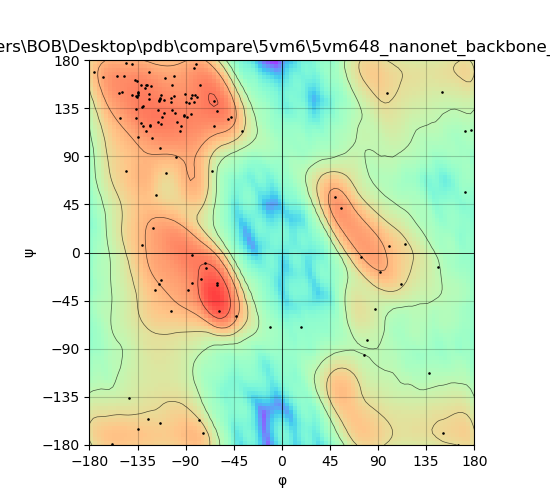

### 6apo_plot_DeepAb.png

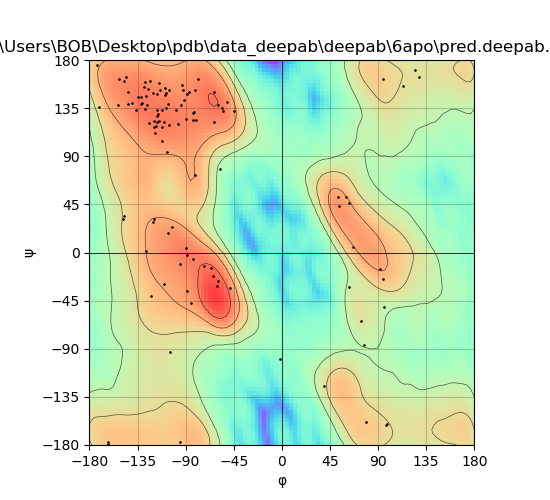

### 6apo_plot_NanoNet.png

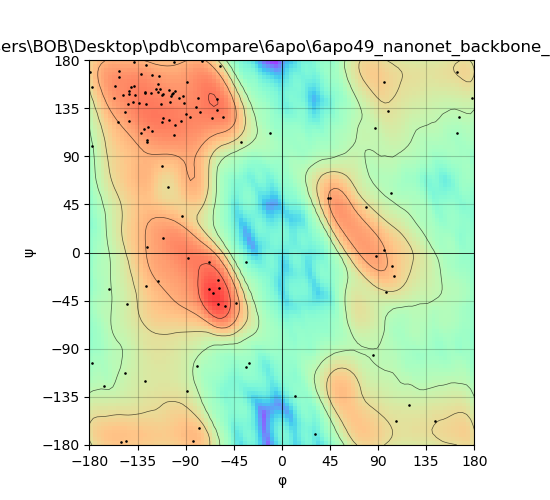

### 6apq_plot_DeepAb.png

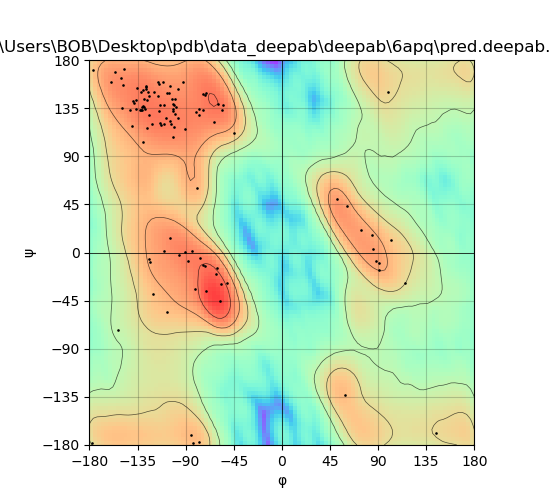

### 6apq_plot_NanoNet.png

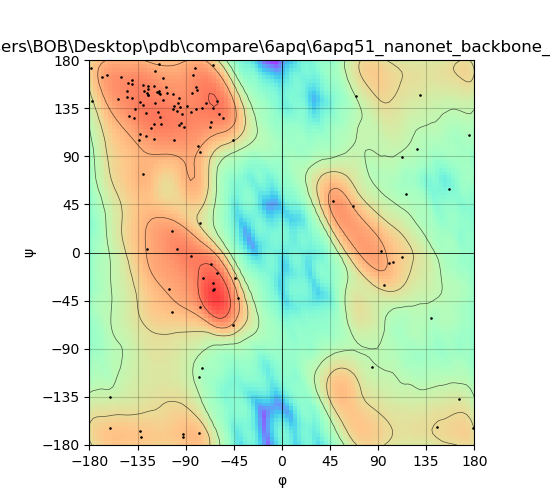

### 6fys_plot_DeepAb.png

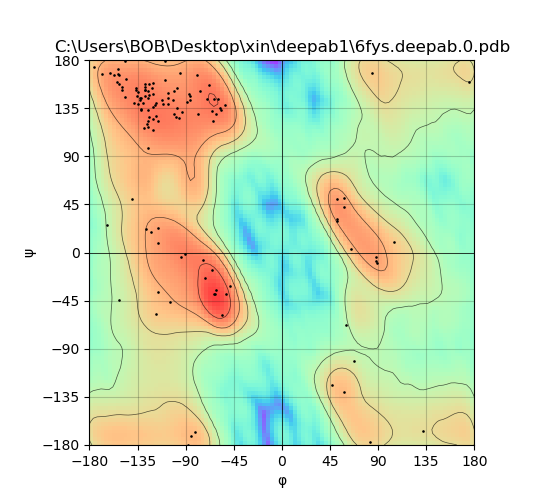

### 6fys_plot_tFold.png

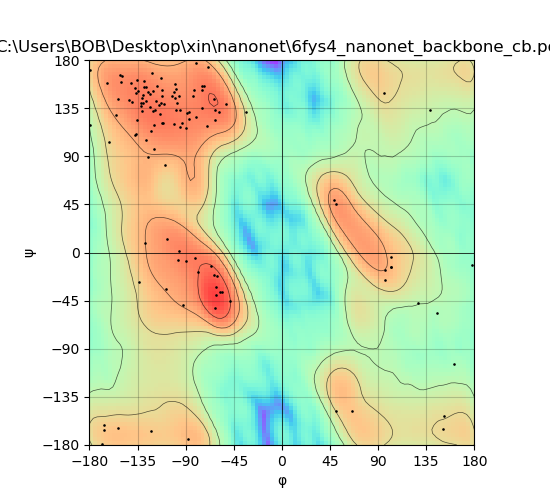

### 6fyt_plot_DeepAb.png

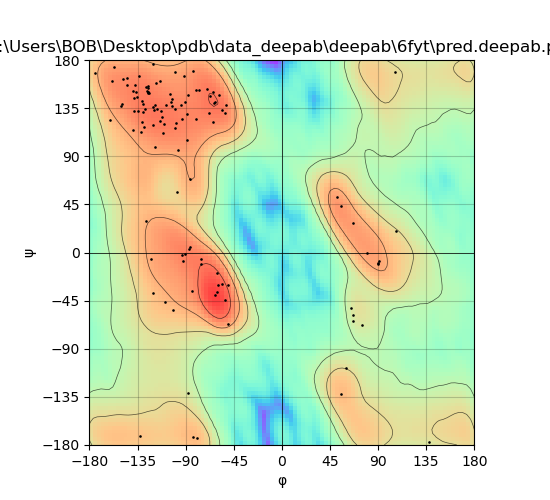

### 6fyt_plot_NanoNet.png

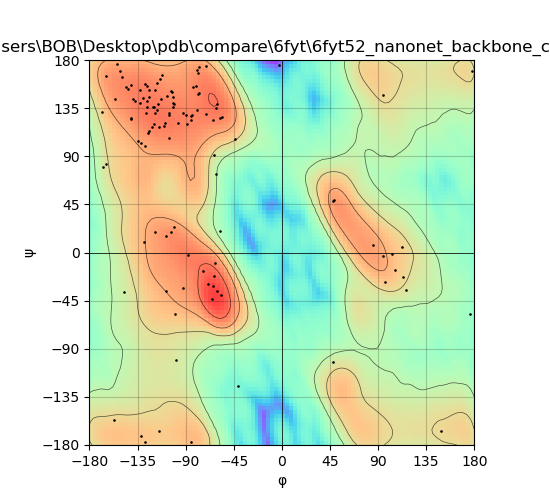

### 6fyw_plot_DeepAb.png

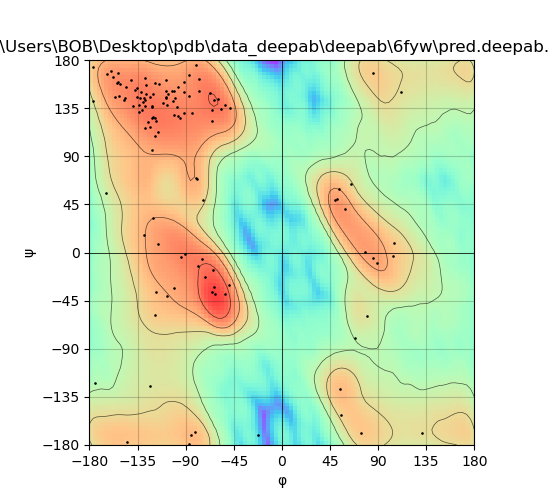

### 6fyw_plot_NanoNet.png

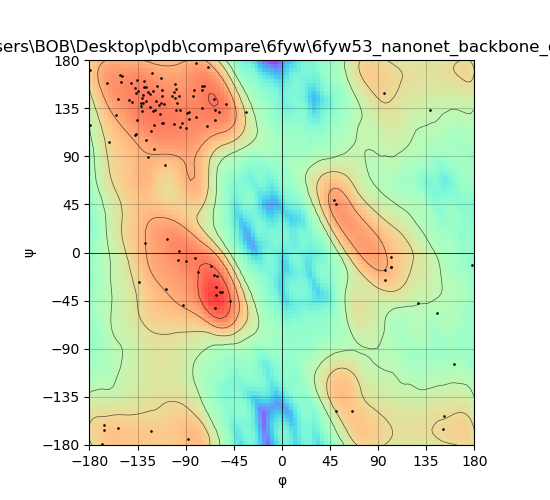

### 6h16_plot_DeepAb.png

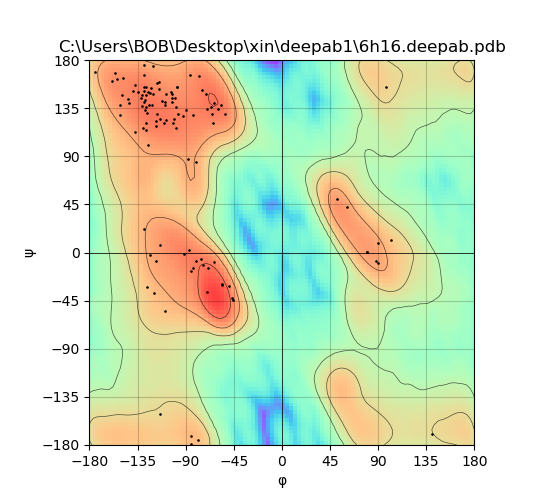

### 6h16_plot_tFold.png

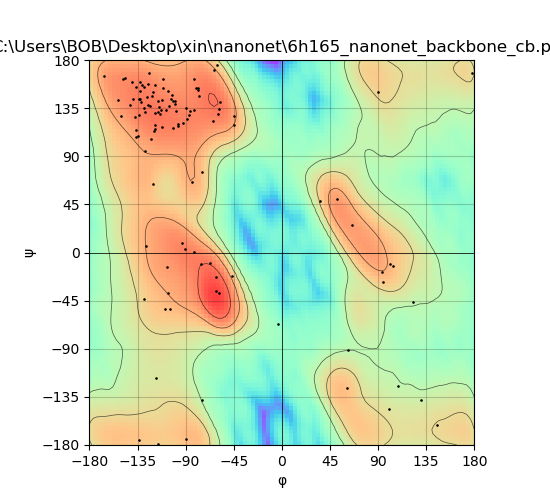

### 6oc8_plot_DeepAb.png

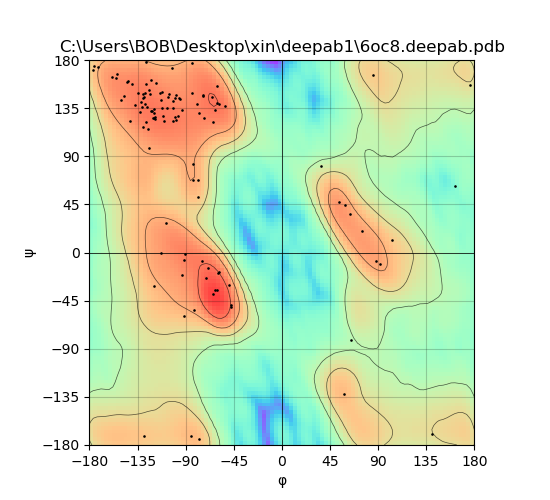

### 6oc8_plot_tFold.png

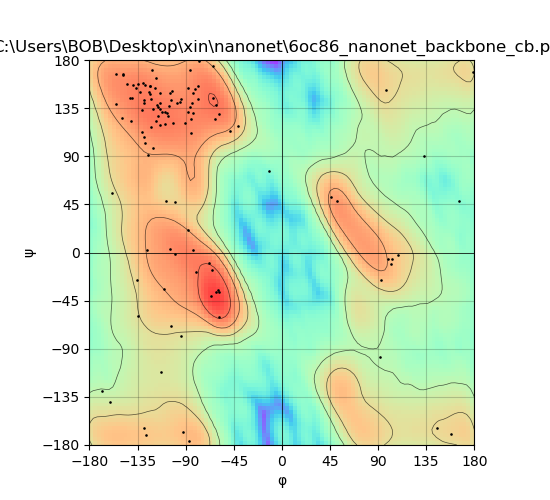

### 6sc5_plot_DeepAb.png

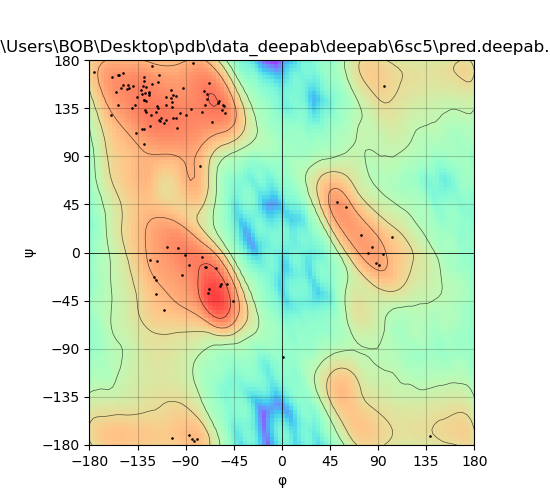

### 6sc5_plot_NanoNet.png

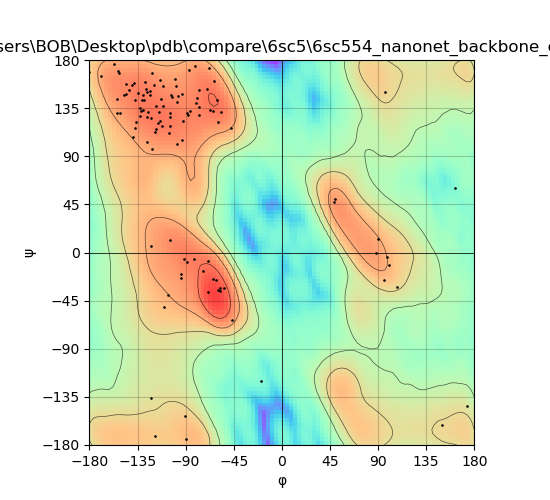

### 6sc8_plot_DeepAb.png

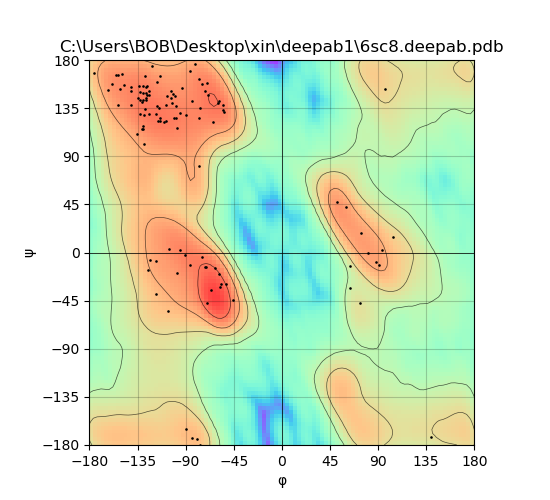

### 6sc8_plot_tFold.png

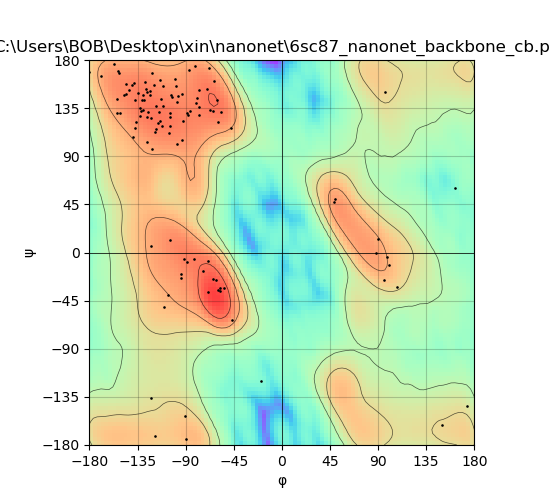

### 6u50_plot_DeepAb.png

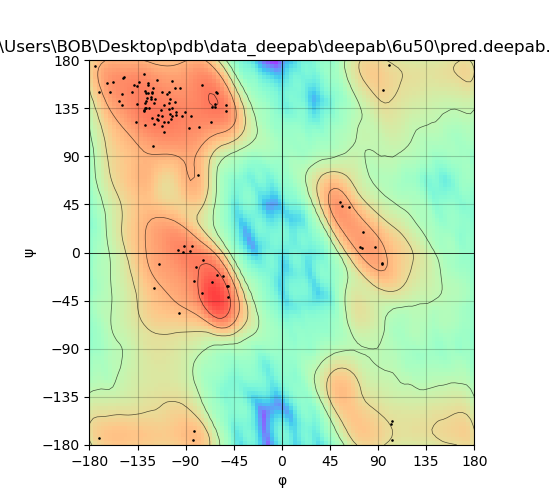

### 6u50_plot_NanoNet.png

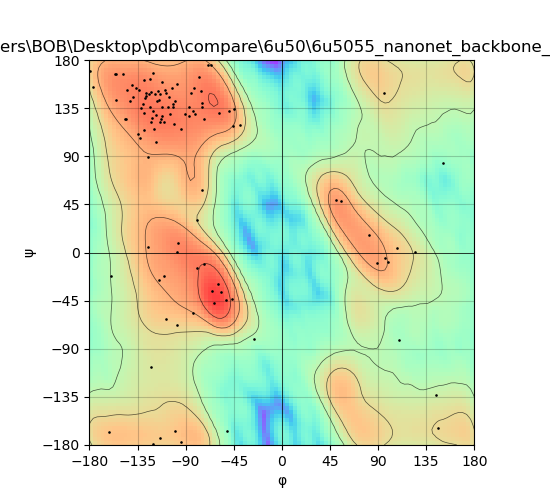

### 6u53_plot_DeepAb.png

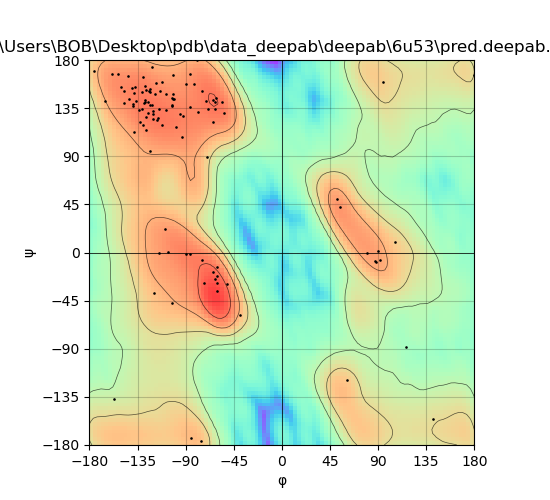

### 6u53_plot_NanoNet.png

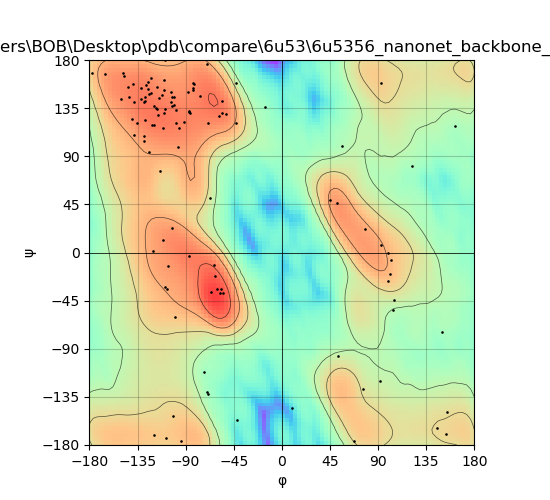

### 6v80_plot_DeepAb.png

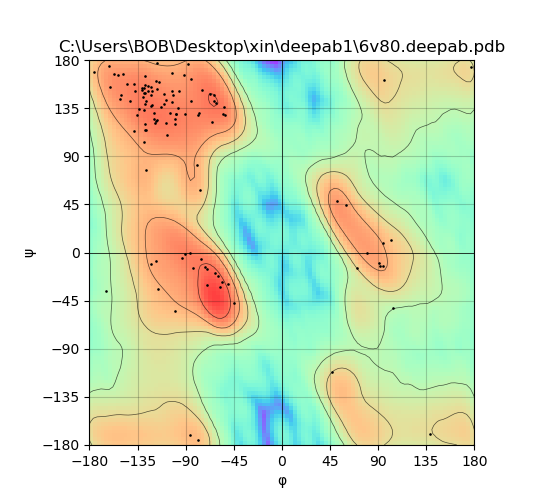

### 6v80_plot_tFold.png

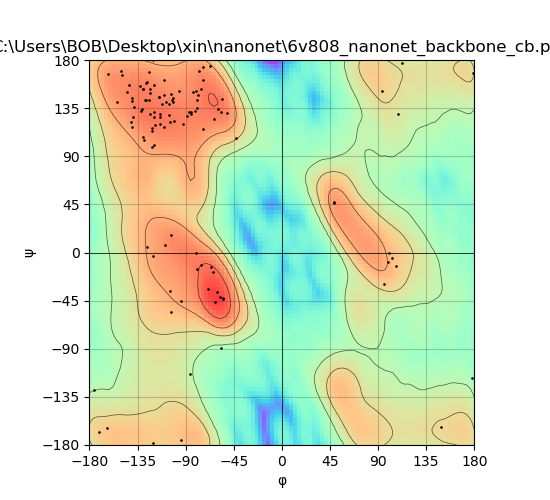

### 7r74_plot_DeepAb.png

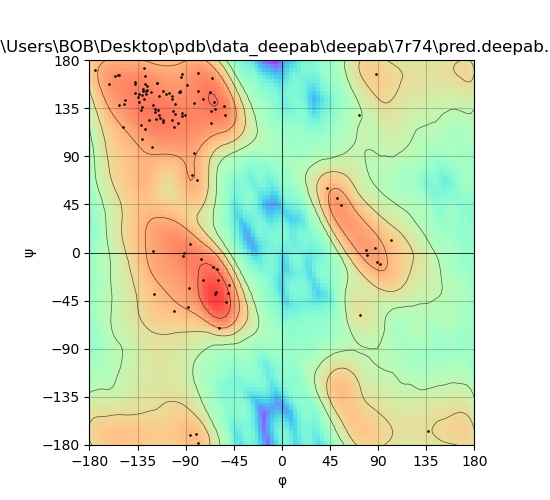

### 7r74_plot_NanoNet.png

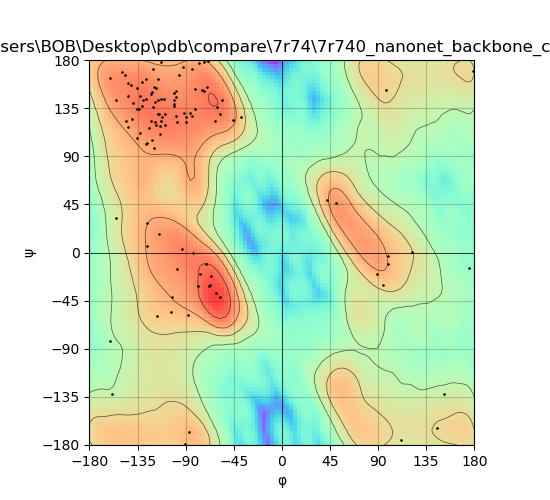

### 7ri1_plot_DeepAb.png

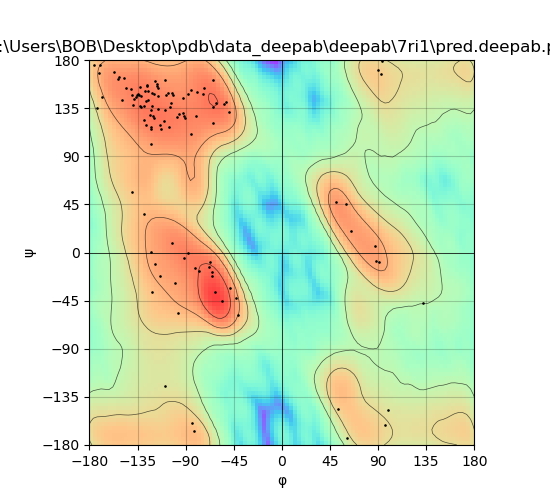

### 7ri1_plot_NanoNet.png

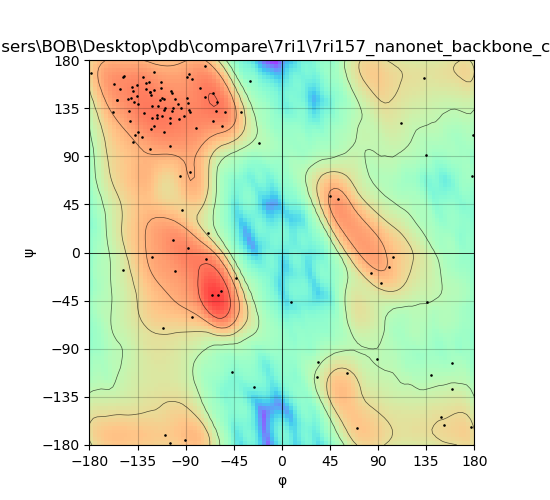
